# Neutrophil subsets play dual roles in tuberculosis by producing inflammasome dependent-IL-1β or suppressing T-cells via PD-L1

**DOI:** 10.1101/2023.11.17.567521

**Authors:** Emilie Doz-Deblauwe, Badreddine Bounab, Florence Carreras, Julia Silveira-Fahel, Sergio C. Oliveira, Mohamed Lamkanfi, Yves Le Vern, Pierre Germon, Julien Pichon, Florent Kempf, Christophe Paget, Aude Remot, Nathalie Winter

**Affiliations:** INRAE, Université de Tours, ISP, F-37380 Nouzilly, France; INSERM, U1100, Centre d’Étude des Pathologies Respiratoires, Tours, France; Faculté de Médecine, Université de Tours, Tours, France; Department of Immunology, University of Sao Paolo, Sao Paulo, Brazil; Department of Biochemistry and Immunology, Federal University of Minas Gerais, Belo Horizonte; Brazil; Laboratory of Medical Immunology, Department of Internal Medicine and Pediatrics, Ghent University, B-9000 Ghent, Belgium

## Abstract

Neutrophils can be beneficial or deleterious during tuberculosis (TB). Based on the expression of MHC-II and programmed death ligand 1 (PD-L1), we distinguished two functionally and transcriptionally distinct neutrophil subsets in the lungs of mice infected with mycobacteria. Inflammatory [MHC-II^-^, PD-L1^lo^] neutrophils produced inflammasome-dependent IL-1β in the lungs in response to virulent mycobacteria and “accelerated” deleterious inflammation, which was highly exacerbated in IFN-γR^-/-^ mice. Regulatory [MHC-II^+^, PD-L1^hi^] neutrophils “brake” inflammation by suppressing T-cell proliferation and IFN-γ production. Such beneficial regulation, which depends on PD-L1, is controlled by IFN-γR signaling in neutrophils. The hypervirulent HN878 strain from the Beijing genotype curbed PD-L1 expression by regulatory neutrophils, abolishing the braking function and driving deleterious hyper-inflammation in the lungs. These findings add a layer of complexity to the roles played by neutrophils in TB and may explain the reactivation of this disease observed in cancer patients treated with anti-PD-L1.

**One Sentence Summary:** Regulatory and inflammatory neutrophil subsets play inverse roles in tuberculosis.

## INTRODUCTION

Tuberculosis (TB) is among the principal causes of death due to infectious diseases in the world. This situation was worsened by the recent burden imposed on healthcare systems by the COVID-19 crisis, which severely affected TB management programs (*1*). Almost all human cases of TB are due to *Mycobacterium tuberculosis* (Mtb). The first laboratory strain sequenced in 1998 by Cole et al. was H37Rv (*2*). It was long believed that genetic diversity among Mtb strains was limited. The recent development of whole genome sequencing uncovered the complex geographical distribution of nine different phylogenetic lineages (L) of Mtb circulating in different regions of the world (*3*). The L2 and L4 strains are the most highly distributed worldwide, with the L2 strain dominating in East-Asia, with high transmission rates. Most experimental TB physiopathology studies have been conducted using the laboratory-adapted L4 strain H37Rv. However, strains from different lineages induce different pathological spectra in humans and animal models (*4*). HN878, the prototype L2 “Beijing” hypervirulent strain, causes an exacerbated immunopathology. However, the immune mechanisms underlying such severe disease are not fully understood.

Following infection with Mtb, most people do not develop immediate signs of disease but may remain latently infected for decades. During this period, a *status quo* between the host and the bacilli involves several immune mechanisms to regulate host defense and inflammation. The role of the programmed death 1/programmed death ligand 1 (PD-1/PD-L1) axis in restricting T-cell function has been recently highlighted. Blockade of these immune check-points has brought considerable progress to cancer treatment in recent years (*5*). However, concerns are now emerging about an increase in active TB cases following such treatment (*5, 6*). Experimentally Mtb-infected PD-1-deficient mice quickly die (*7*) due to the detrimental overproduction of pathogenic IFN-γ by CD4^+^ T cells in the lung parenchyma (*8*). Mtb-infected rhesus macaques treated with anti-PD-1 develop exacerbated disease, which is linked to caspase-1 activation (*9*). Inherited PD-1 deficiency in humans is linked to decreased self-tolerance and anti-mycobacterial immunity (*10*).

The hallmark of TB is the formation of granulomas in the lung; in these organized pluricellular structures, a delicate balance between the containment of Mtb replication and host inflammation takes place. The fate of Mtb, from eradication to active multiplication, may vary depending on the granuloma microenvironment, where multiple immune mechanisms are at play to maintain or disrupt immunoregulation (*11*). Among innate cells, neutrophils play dual roles in TB (*12*). At early stages, they halt Mtb infection and shape early formation of the TB granuloma (*13, 14*). At later stages, their highly destructive arsenal is critical for TB reactivation; they represent the first expectorated cells of active TB patients (*15*). In the mouse, we have shown that neutrophils reach the lungs in two waves during the establishment of the immune response, with the adaptive wave playing no role in Mtb growth restriction (*16*). There is now extensive evidence that neutrophils represent a heterogeneous and plastic cellular compartment (*17*). Some neutrophils are endowed with classical phagocytic and pathogen-killing functions, whereas others are able to cross talk with a variety of immune cells, taking full part in the adaptive immune response (*12*). Although the dual roles played by neutrophils are recognized in TB, the possibility that distinct subsets play divergent roles is still underexplored (*18*). In this context, we have recently characterized a subset of regulatory neutrophils that can be functionally distinguished from classic neutrophils in healthy cattle and mice by their ability to suppress T-cell proliferation (*19*).

IL-1β is a cornerstone cytokine in TB. It is essential for constraining Mtb infection in the early stages, as unequivocally demonstrated in mouse models, but may also become deleterious at later stages of the full-blown adaptive immune response. Cross-regulatory pathways of IL-1β production during TB include that of type I IFN, which directly downregulates pro-IL-1β gene transcription (*20*). Bioactive IL-1β needs to be processed from immature pro-IL-1β via inflammasome assembly, to which macrophages (MPs) are the major contributor. *In vitro*, in response to Mtb infection, bone marrow-derived MPs assemble the NLRP3 inflammasome and activate caspase-1 to trigger canonical inflammasome activation and the release of mature IL-1β (*21*). Beyond MPs, recent studies suggest a role for NLRP3 inflammasome-dependent IL-1β production by neutrophils *in vivo* (*22*). However, the contribution of neutrophils to IL-1β production during TB appears to be much less than that of MPs (*23*) and it is assumed that caspase-1-independent mechanisms account for pro-IL-1β cleavage by these cells (*24*).

As neutrophils shape the fate and full development of granulomas during TB disease, we revisited the role of these heterogeneous plastic cells (*17, 19*) during mycobacterial infection. We compared the recruitment and functions of neutrophil subsets during infection with the avirulent live vaccine BCG and the two virulent Mtb strains, H37Rv (L4, lab-adapted) and HN878 (L2, Beijing prototype). We used the IFN-γR^-/-^ mouse model, in which extensive neutrophil-driven inflammation was described (*25*) before distinct subsets were known. We also analyzed the potential for inflammasome-dependent mature IL-1β production by neutrophils *in vitro*, as well as *in vivo*, taking advantage of a new mouse model in which caspase-1-dependent IL-1β secretion is specifically abrogated in neutrophils (*26*). We provide evidence that distinct subsets play opposite roles in TB physiopathology by contributing to IL-1β-driven inflammation in the lungs or regulating neutrophilia via the immune checkpoint inhibitor PD-L1.

## RESULTS

### The neutrophil NLRP3 inflammasome contributes to IL-1**β** production during mycobacterial infection

We infected neutrophils from mouse bone marrow with the avirulent vaccine strain BCG and the virulent H37Rv and HN878 Mtb strains. Different MOIs for the BCG (10:1) virulent Mtb (1:1) were used to preserve neutrophil viability. This induced comparable release of TNF (Fig. S1A). Neutrophils also released mature IL-1β in response to infection by all strains (Fig. 1A), albeit to a lesser extent than following LPS plus nigericin stimulation. We next prepared neutrophils from the bone marrow of various genetically deficient mice to test the role of the inflammasome. IL-1β secretion by mycobacteria-infected or LPS/nigericin stimulated neutrophils from *Nlrp3*^-/-^ (Fig. 1B), *Csp1/11* ^-/-^ (Fig. 1B), and *Gsdmd*^-/-^ (Fig. 1C) mice was severely impaired relative to that of WT neutrophils. This was not due to activation issues, as these genetically deficient neutrophils released similar levels of TNF (Fig. S1B). Canonical assembly of the inflammasome and pyroptosis appear to be involved in the IL-1β maturation process in neutrophils. This was confirmed with neutrophils from *Csp11*^-/-^ mice, which secreted similar levels of mature IL-1β as WT mice in response to BCG infection (Fig. S1C). Of note, neutrophils from *Aim2*^-/-^ or WT mice produced similar levels of mature IL-1β (Fig. 1C). We observed the cleavage of pro-IL1β into mature IL-1β of 17kDa in neutrophils infected with BCG (MOI 20:1) by western blotting (Fig. 1D), confirming inflammasome assembly. MPs secreted more mature IL-1β into the supernatant than neutrophils (Fig. 1E), regardless of the stimulus. On a cell-to-cell basis, MPs secreted 47 times more mature IL-1β than neutrophils after infection with virulent Mtb H37Rv and 35 times more than after BCG stimulation (Fig. 1E). We assessed the contribution of neutrophils to IL-1β production *in vivo* by infecting mice with virulent Mtb H37Rv and injecting the anti-Ly6G antibody at the onset of recruitment of the second wave of neutrophils, i.e., between day 17 and 21 (*16*). This treatment markedly reduced the number of neutrophils in the lungs (Fig. 2A and Fig. S2A for gating strategy). Lesions were more extensive in anti-Ly-6G than isotype treated mice (Fig. 2B), with a two-fold greater total lung surface occupied by the lesions (Fig. 2C). Production of IL-1β in the lung tissue of anti-Ly-6G treated mice was 2.3-fold less than that in the lung tissue of mice injected with the isotype control antibody (Fig. 2D). These data thus confirm the role of neutrophils in the formation of lung lesions during Mtb infection (*12*) and indicate their direct participation in IL-1β production *in vivo*. We confirmed this using the recently obtained *MRP8^Cre+^Csp1^flox^* mouse strain (*26*), in which IL-1β production is specifically abolished in neutrophils. We first validated this tool *in vitro* using purified neutrophils and bone marrow-derived MPs from *MRP8^Cre+^Csp^flox^* mice and their *MRP8^WT^Csp1^flox^* littermates stimulated with LPS and nigericin. As expected, neutrophils from *MRP8^Cre+^Csp1^flox^*mice did not produce IL-1β, whereas *MRP8^WT^Csp1^flox^*neutrophils did (Fig. 2E). In addition, MPs from *MRP8^Cre+^Csp1^flox^*and *MRP8^WT^Csp1^flox^* mice equally produced IL-1β, as expected (Fig. 2F). Next, we intranasally infected *MRP8^Cre+^Csp1^flox^* mice and their *MRP8^WT^Csp1^flox^* littermates with Mtb H37Rv and observed significantly lower IL-1β production in the lungs of the *MRP8^Cre+^Csp1^flox^* than those of the *MRP8^WT^Csp1^flox^*animals (Fig. 2G). This result confirms the direct participation of neutrophils in IL-1β production after inflammasome assembly in the lungs in response to Mtb infection.

**Fig. 1.**
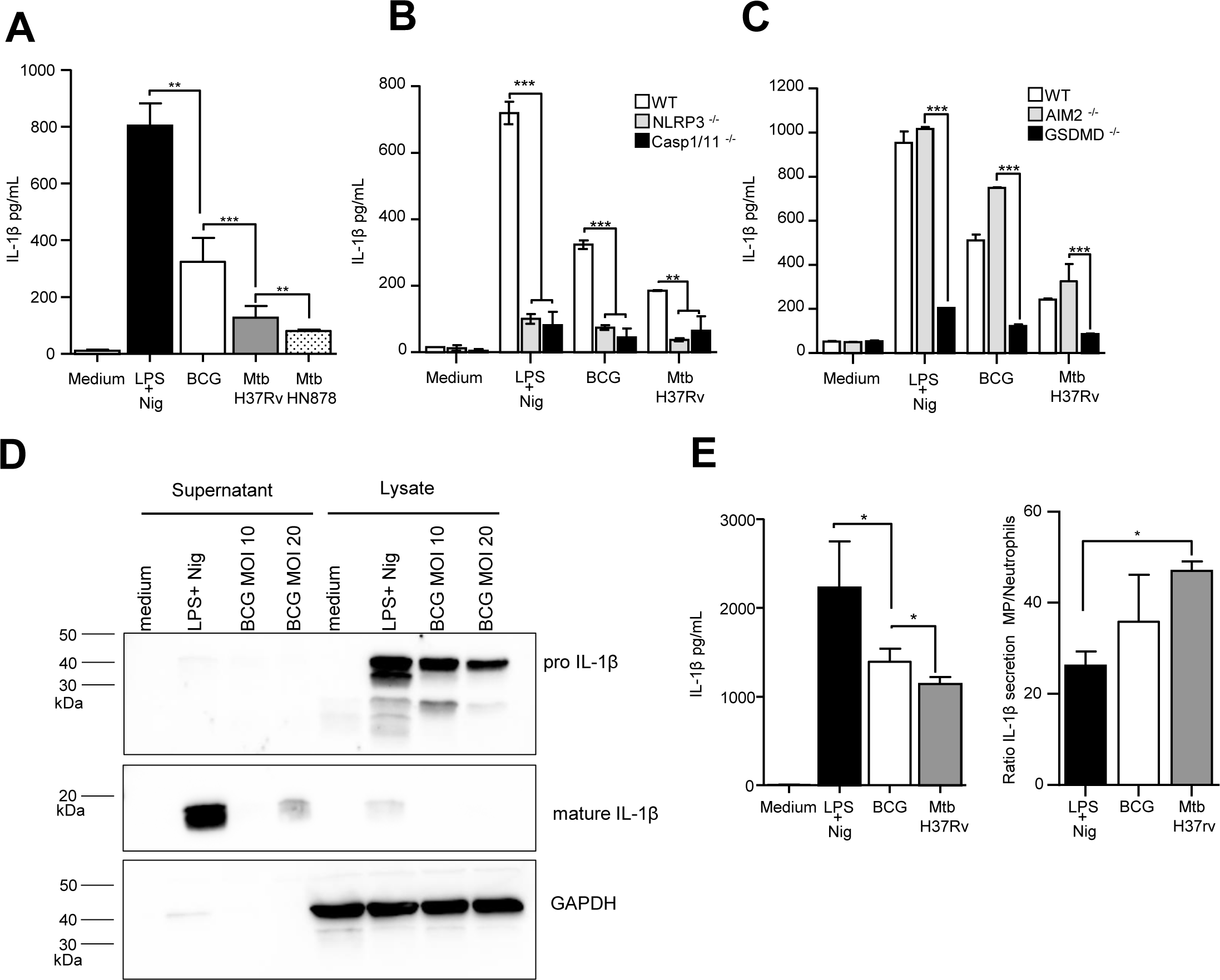
The neutrophil NLRP3 inflammasome contributes to IL-1β production after mycobacterial infection. Mature IL-1β produced by WT **(A)**, *Nlrp3*^-/-^, *Csp1/11*^-/-^ **(B)**, *Aim2*^-/-^, and *Gsdmd*^-/-^ **(C)** bone marrow neutrophils was determined by ELISA after overnight stimulation with LPS/nigericin or infection with BCG (MOI 10) or Mtb (H37Rv or HN878, MOI 1). **(D)** Immunoblotting of pro-IL-1β, mature IL-1β, and GAPDH in supernatants and lysates from bone marrow neutrophils infected for 5 h with BCG (MOI 10 or 20) or stimulated with LPS/nigericin. **(E)** Mature IL-1β produced by WT bone marrow-derived MPs was determined by ELISA after overnight stimulation with LPS/nigericin or infection with BCG (MOI 10) or Mtb H37Rv (MOI 1). (A) Pooled data from three independent experiments, n = 6 mice; (B) data are representative of three independent experiments, n = 3; (C) data are representative of two independent experiments, n = 3; (D) data are representative of two independent experiments, n = pool of 10 mice; (E) pooled data from two independent experiments, n = 4. Graphs show medians with ranges. ∗*p* < 0.05, ∗∗*p* < 0.01, and ∗∗∗*p* < 0.001 001 by the Mann-Whitney test (A, E) and two-way ANOVA with a Bonferroni post-test (B, C).

**Fig. 2.**
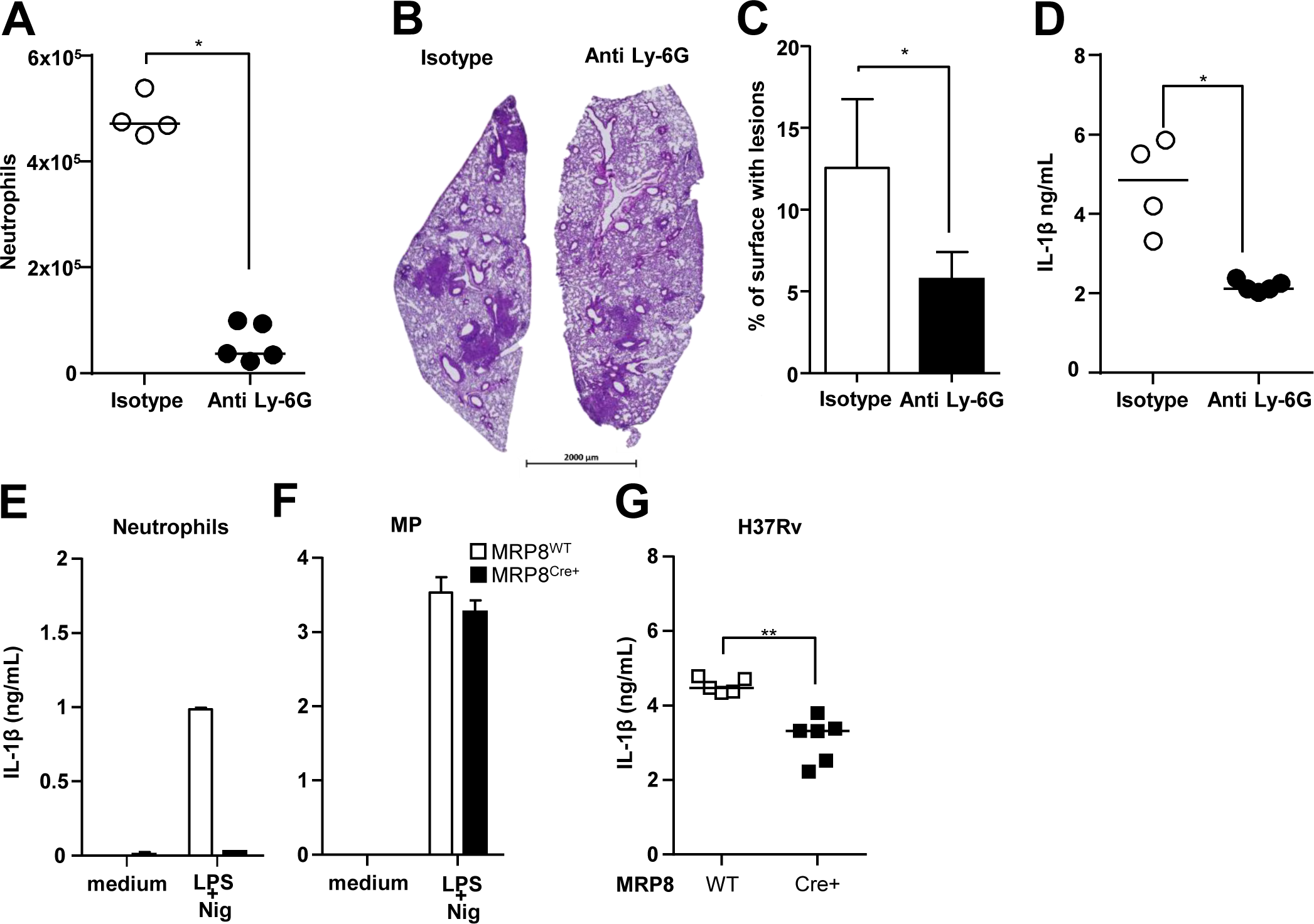
Neutrophils directly contribute to IL-1β production in the lungs during H37Rv infection. **(A-C)** C57BL/6 mice were infected with 10^3^ CFUs of Mtb H37Rv and neutrophils depleted by intraperitoneal administration of anti-Ly6G or isotype control antibody on days 15, 17, and 19. Lungs were harvested on day 21 for analysis. **(A)** Total lung neutrophils were identified by flow cytometry as CD11b^+^ Ly6G^+^ Ly6C^+^ cells (see Fig S2a for gating strategy). **(B)** Representative image of hematoxylin/eosin staining of lung sections. (**C**) Total lung surface occupied by lesions in lung sections from anti-Ly-6G or isotype control antibody-treated mice. **(D)** IL-1β production in lung tissue from C57BL/6 mice or in the supernatants after overnight stimulation with LPS/nigericin of bone marrow neutrophils (**E)** or MPs **(F)** from *MRP8^Cre+^Csp1^flox^* or *MRP8^WT^Csp1^flox^* mice measured by ELISA. **(G)** IL-1β production in lung tissue from *MRP8^Cre+^Csp1^flox^* or *MRP8^WT^Csp1^flox^* mice 21 days post-infection with H37Rv measured by ELISA. (A-D) One experiment, n = 4-5 mice per group; (E, F) two independent experiments, n = 4 mice; (G) data representative of two independent experiments, n = 5-6 per group. Medians with ranges (C, E, F) and individual data points with medians for A, D, and G). ∗*p* < 0.05, ∗∗*p* < 0.01 by the Mann-Whitney test (A-D) and two-way ANOVA with a Bonferroni post-test (E-F).

### Mycobacteria attract inflammatory and regulatory neutrophil subsets to the lung

We recently discovered that [Ly-6G^+^, MHC-II^-^, PD-L1^lo^] neutrophils, akin to classic neutrophils, and [Ly-6G^+^, MHC-II^+^, PD-L1^hi^] regulatory neutrophils circulate in blood as two functionally different subsets at steady state in healthy mice and cattle. Only regulatory neutrophils are able to suppress T-cell proliferation (*19*). Thus, we first assessed the recruitment of these two neutrophil subsets to the lungs following intranasal infection with 5×10^6^ CFUs of avirulent BCG. Surface MHC-II was used to discriminate between classic and regulatory neutrophils by flow cytometry (Fig. S2A). As previously observed (*16*), total [CD45^+^, CD11b^hi^; Ly-6G^hi,^ Ly-6C^+^] neutrophils peaked in the lungs 21 days following BCG infection (Fig. 3A), together with T cells. This cell population was composed of a balanced mix of [Ly-6G^+^, MHC-II^-^] classic neutrophils and [Ly-6G^+^, MHC-II^+^] regulatory neutrophils (Fig. 3B), which showed similar morphology (Fig. S2B). PD-L1 also clearly distinguished [Ly-6G^+^, MHC-II^-^, PD-L1^lo^] from [Ly-6G^+^, MHC-II^+^, PD-L1^hi^] neutrophils (Fig. S2A). Overall, 90% of [Ly-6G^+^, MHC-II^+^] regulatory neutrophils were PD-L1^hi^ and 10% of [Ly-6G^+^, MHC-II^-^] classic neutrophils were PD-L1^lo^ (Fig. 3C). Moreover, the MFI of [Ly-6G^+^, MHC-II^+^, PD-L1^hi^] neutrophils was 26 times higher than that of [Ly-6G^+^, MHC-II^-^, PD-L1^lo^] neutrophils (Fig. 3D). We then performed single-cell RNAseq analysis of the total Ly-6G^+^ neutrophil population purified from the lungs 21 days after BCG infection and observed distinct transcriptional profiles (Fig 3E). SEURAT software classified RNA expression into clusters numbered 0 to 10 (Fig 3E, first panel) that formed two main groups: clusters 0, 2, 3, 4, and 5 formed part of one pool (Fig. 3E, right pool), whereas clusters 1, 6, 7, 8, 9, and 10 formed another (Fig. 3E, left pool). Of note the SingleR software, trained on the Immunologic Genome Project database of mRNA profiles, identified cells from the right pool as “neutrophils”, whereas cells from the left pool were identified as “monocytes/macrophages”, probably due to the expression of genes such as *Mhc-II* and *Cd274* (encoding PD-L1). Certain genes, such as *Itgam* (Fig. 3E), were similarly expressed in clusters from the two pools, in agreement with the neutrophil signature. However, differential expression of the *H2-Eb1* and *H2-Ab1* genes from the MHC-II complex or *Il1b* and *Il1r2* inflammatory genes clearly segregated between the two pools (Fig. 3E). We confirmed the differential gene expression between classic and regulatory neutrophils by performing qRT-PCR targeting genes involved in general neutrophil-driven inflammation, as well as pro-IL-1β synthesis and inflammasome assembly. *Mhc-II* genes were expressed in regulatory neutrophils only (Fig. 3F). We observed full differential clustering of the two subsets based on the expression of *Il1b*, *Ilr2*, *Mmp9*, *Ifngr*, *Il18rap,* and *S100a9*; the expression of all these genes was higher in [Ly-6G^+^, MHC-II^-^, PD-L1^lo^] classic neutrophils than [Ly-6G^+^, MHC-II^+^, PD-L1^hi^] regulatory neutrophils (Fig. 3F). We separated the two subsets from H37Rv-infected mice using magnetic beads and observed higher *Il1b* and *Ilr2* gene expression by classic than regulatory neutrophils (Fig. S2C). We then analyzed intracellular mature IL-1β production in the lungs *ex vivo* by flow cytometry three weeks following H37Rv infection (Fig. S2D). The MFI for IL-1β was 25 times higher in [Ly-6G^+^, MHC-II^-^] classic than [Ly-6G^+^, MHC-II+] regulatory neutrophils (Fig. 3G). Based on these data, we considered classic neutrophils to be “inflammatory” during mycobacterial infection, as documented both by their transcriptional profile and their ability to produce mature IL-1β *in vivo*.

**Fig. 3.**
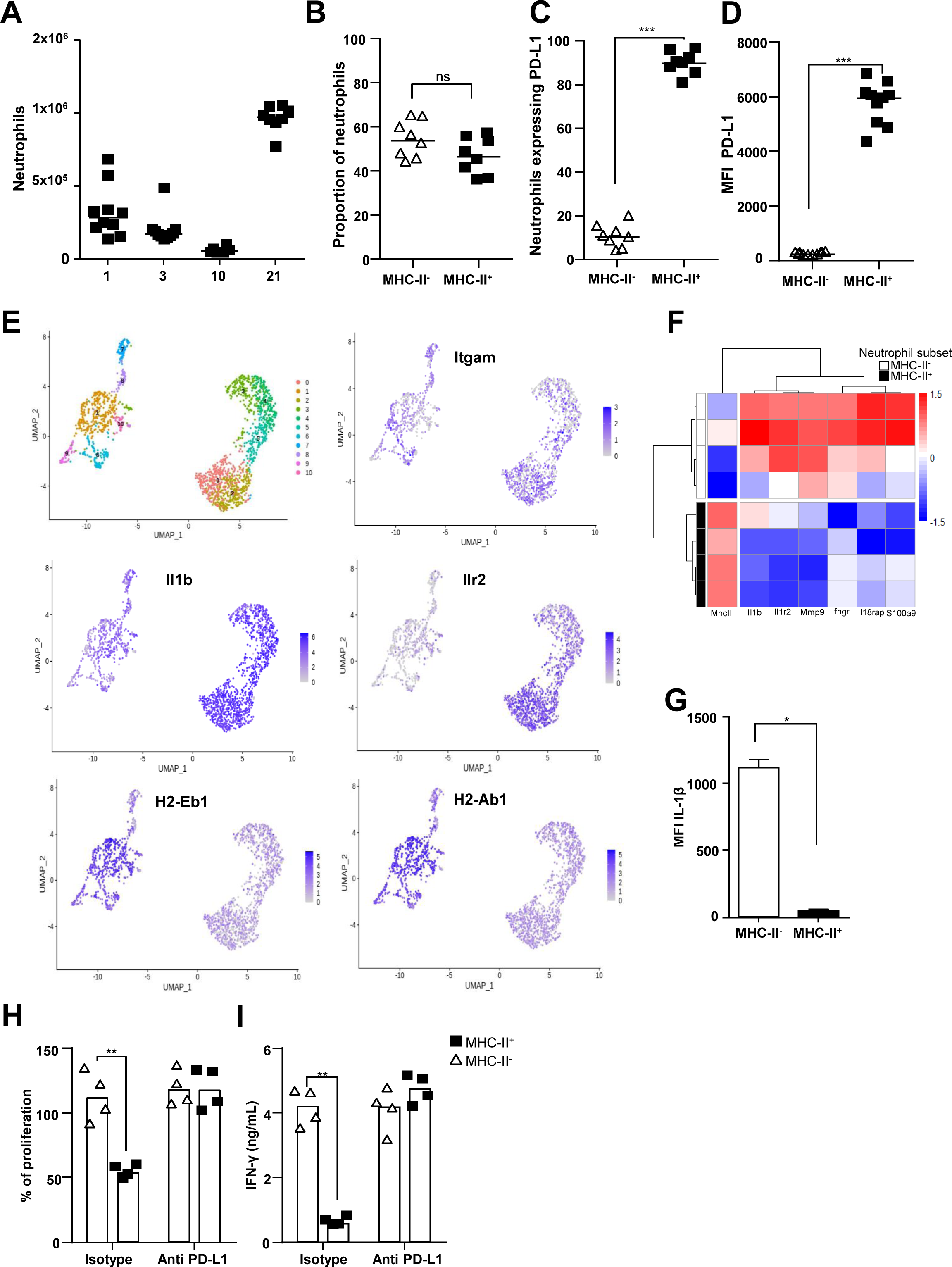
Mycobacteria attract inflammatory and regulatory neutrophil subsets to the lung. **(A-E)** C57BL/6 WT mice were infected with BCG and the lungs processed for following analysis. (**A)** Kinetics of total lung neutrophils recruited to the lungs at days 1, 3, 10, and 21 post-infection assessed by flow cytometry (Fig. S2A for gating strategy). **(B-F)** Lung neutrophils were further characterized on day 21 post-infection. **(B)** Proportion of MHC-II^+^ and MHC-II^-^ neutrophils among total lung neutrophils. **(C)** Percentage of PD-L1 expression among neutrophils in each subset. (**D**) PD-L1 mean fluorescence intensity in each subset. **(E)** Single-cell RNAseq analysis of Ly6G^+^ neutrophils purified from a pool of lung cells from 10 mice. Identification of 11 cell clusters using the SEURAT package on Uniformed Manifold Approximation and Projections (UMAP). Each dot represents one cell. Visualization of *Itgam*, *Il-1b*, *Il-1R2*, *H2-Eb1*, and *H2-Ab1* gene expression in the clusters as analyzed by the SEURAT package. **(F)** Heatmap representation of differential gene expression between MHC-II^-^ and MHC-II^+^ neutrophils. **(G)** C57BL/6 WT mice were infected with Mtb H37Rv and mean fluorescence intensity of intracellular IL-1β measured by flow cytometry in both MHC-II^-^ and MHC-II^+^ lung neutrophil subsets on day 21. **(H)** MHC-II^-^ and MHC-II^+^ neutrophil subsets were enriched by magnetic beads from the lungs of H37Rv-infected mice on day 21 and mixed with OT-II cells. The percentage of OT-II splenocyte proliferation in the presence of each neutrophil subset was calculated based on proliferation with the Ova peptide only. Neutrophils were treated 1 h before incubation with anti PD-L1 Ab (atezolizumab) or an isotype control. (**I**) IFN-γ production in supernatants of OT-II splenocytes measured by ELISA. (A) Pooled data from two independent experiments (n = 8-10 per group); (B-D) pooled data from two independent experiments (n = 8 per group); (E) one experiment, pool of 10 mice; (F) four cell sorting experiments were performed from a pool of five infected animals each time; (G-I) pooled data from two independent experiments (n = 4). Data are presented as individual data points and medians. ∗*p* < 0.05, ∗∗*p* < 0.01, and ∗∗∗*p* < 0.001 by the Mann-Whitney test (A-D, G) and two-way ANOVA with a Bonferroni post-test (H-I).

As the immune inhibitory checkpoint PD-L1 is involved in T-cell suppression (*27*), we assessed its role in lung regulatory neutrophils recruited in response to mycobacterial infection. We separated regulatory from inflammatory neutrophils from the lungs of BCG- or Mtb H37Rv-infected mice on day 21 and tested their suppressive function *ex vivo* on splenocytes from OT-II mice (Fig S2E and (*19*). Only the [Ly-6G^+^, MHC-II^+^, PD-L1^hi^] regulatory neutrophils were able to decrease OT-II cell proliferation (by 50%, Fig. 3H) and IFN-γ production (by 87%, Fig. 3I). We observed similar levels of T-cell suppression by lung regulatory neutrophils obtained from BCG-(Fig. S2F) or Mtb H37Rv-infected mice (Fig. 3H). Moreover, addition of the anti-PD-L1 antibody atezolizumab (*28*) to the wells with regulatory neutrophils obtained from H37Rv-(Fig. 3H) or BCG-(Fig. S2F) infected mice fully restored proliferation and IFN-γ production by OT-II cells. Thus, only regulatory neutrophils were able to dampen T-cell function and PD-L1 played a major role in this effect.

### The two neutrophil subsets are modulated by *M. tuberculosis* virulence

We next sought information on the role of the neutrophil subsets in TB physiopathology by comparing lung infection by H37Rv and HN878 on day 21. The number of bacilli in the lungs was 1.1 log_10_ higher (Fig. 4A) and the lesions (Fig. 4B) occupied 4.7 times more lung surface (Fig. 4C) in HN878 than H37Rv-infected animals, in agreement with the hypervirulence of the Beijing strains (*29, 30*). However, all mice were clinically stable until the end of our study, i.e., day 21 (data not shown). An analysis of differentially expressed genes (Fig. 4D) showed higher expression of *Sting1, Irf3*, *Ifnar1*, and *Ifnar2* from the type I IFN pathway in H37Rv than HN878 infected animals. On the contrary, expression of the neutrophil marker genes *S100a8* and *S100a9* was higher in the lungs of HN878-than those of H37Rv-infected mice. However, the genes involved in inflammasome assembly and the IL-1β production pathway were not distinctly induced by the two virulent Mtb strains. At the protein level, TNF, IFN-γ, and IL-1β levels were higher in the lungs of HN878 than those of H37Rv-infected mice (Fig. 4E), in agreement with the strong inflammatory profile of the strain. CXCL-10 levels, a promising biomarker of Mtb infection (*31*), immunohistochemistry and the hypervirulence of HN878 linked to strong neutrophilia (*32*). However, although the total neutrophil influx was composed of 59% inflammatory and 41% regulatory neutrophils after H37Rv infection, HN878 resulted in an opposite balance of 71% regulatory and 29% inflammatory neutrophils (Fig. 4G). Moreover, infection with HN878 induced a mean MFI for PD-L1 on lung regulatory neutrophils that was 3.8 times lower than that for those of H37Rv-infected animals (Fig. 4H).

**Fig. 4.**
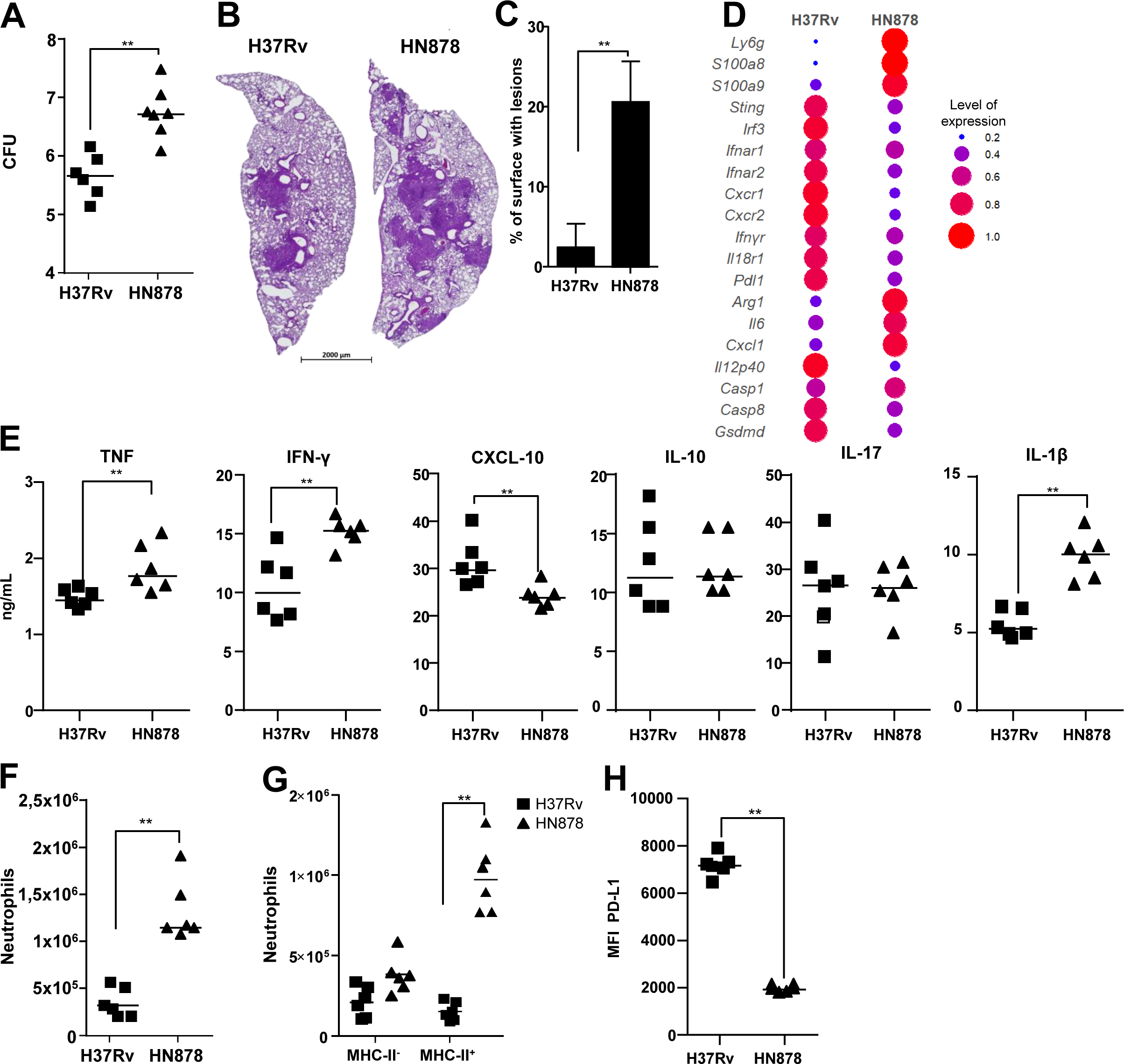
The two neutrophil subsets are modulated by *M. tuberculosis* virulence. **(A-G)** All data (n = 6 mice per group, two independent experiments) were obtained for the lungs of C57BL/6 WT mice at 21 days post-infection with Mtb H37Rv or HN878. **(A)** Number of Mtb CFUs in the lungs. **(B)** Representative image of hematoxylin/eosin staining of lung sections for each infected group and **(C)** the mean percentage of lung surface occupied by lesions. **(D)** Differential gene expression between the two infected groups of a panel of 48 genes normalized against uninfected controls. The dot plot represents the normalized expression of significantly deregulated genes, expressed as the normalized rate to compare the two groups. Data are presented as the mean of n = 4 mice per group from one experiment. **(E)** Cytokine production as analyzed by ELISA in lung tissue homogenates. **(F)** Total neutrophils, **(G)** neutrophil subsets, and **(H)** PD-L1 surface expression by MHCII^+^ neutrophils from the two groups measured by flow cytometry. Data are presented as individual data points and medians (B, median with range). (A-H) Two independent experiments, n = 6 mice per group. ∗*p* < 0.05, ∗∗*p* < 0.01, and ∗∗∗*p* < 0.001 by the Mann-Whitney test (A-E, F, H) and two-way ANOVA with a Bonferroni post-test (G).

### Caspase-dependent production of IL-1**β** by inflammatory neutrophils sustains lung inflammation

Inflammatory neutrophils produced mature IL-1β after NLRP3 inflammasome assembly *in vivo*. We next addressed their contribution to IL-1β-mediated physiopathology in *MRP8^Cre+^Csp1^flox^* mice. We intranasally infected *MRP8^Cre+^Csp1^flox^* mice and *MRP8^WT^Csp1^flox^*littermates with avirulent BCG and the two virulent H37Rv and HN878 Mtb strains and analyzed their response in the lungs three weeks later. We first observed that, in response to BCG, IL-1β levels in whole lung tissue homogenates were low and comparable in *MRP8^Cre+^Csp1^flox^* and *MRP8^WT^Csp1^flox^*animals (Fig. S3A). On the contrary, IL-1β production by the lungs was lower in *MRP8^Cre+^Csp1^flox^*than *MRP8^WT^Csp1^flox^* animals in response to the two virulent Mtb strains. While the response to H37Rv in terms of the amount of IL-1β in the lungs of *MRP8^Cre+^Csp1^flox^*mice was only 30% lower than that of *MRP8^WT^Csp1^flox^*control mice, it was reduced by 64% in response to HN878 (Fig. 5A). Neutrophil-derived IL-1β had no impact on the number of CFUs after infection with BCG (Fig. S3B), H37Rv, or HN878 (Fig 5B) at the time point examined. We next examined the differential lung gene expression profile between *MRP8^Cre+^Csp1^flox^*and *MRP8^WT^Csp1^flox^* mice after infection with BCG (Fig. S3C) or the virulent Mtb strains (Fig. 5C) on day 21. In response to the three strains, *Cxcl5*, a critical gene for neutrophil recruitment to the lungs (*16, 33*), as well as *Cxcl10*, were more highly expressed when inflammatory neutrophils were able to produce IL-1β than when they were defective. In addition, HN878 induced higher transcription of *Il-10* and *Cxcr1* when inflammatory neutrophils were defective for IL-1β production.

**Fig. 5.**
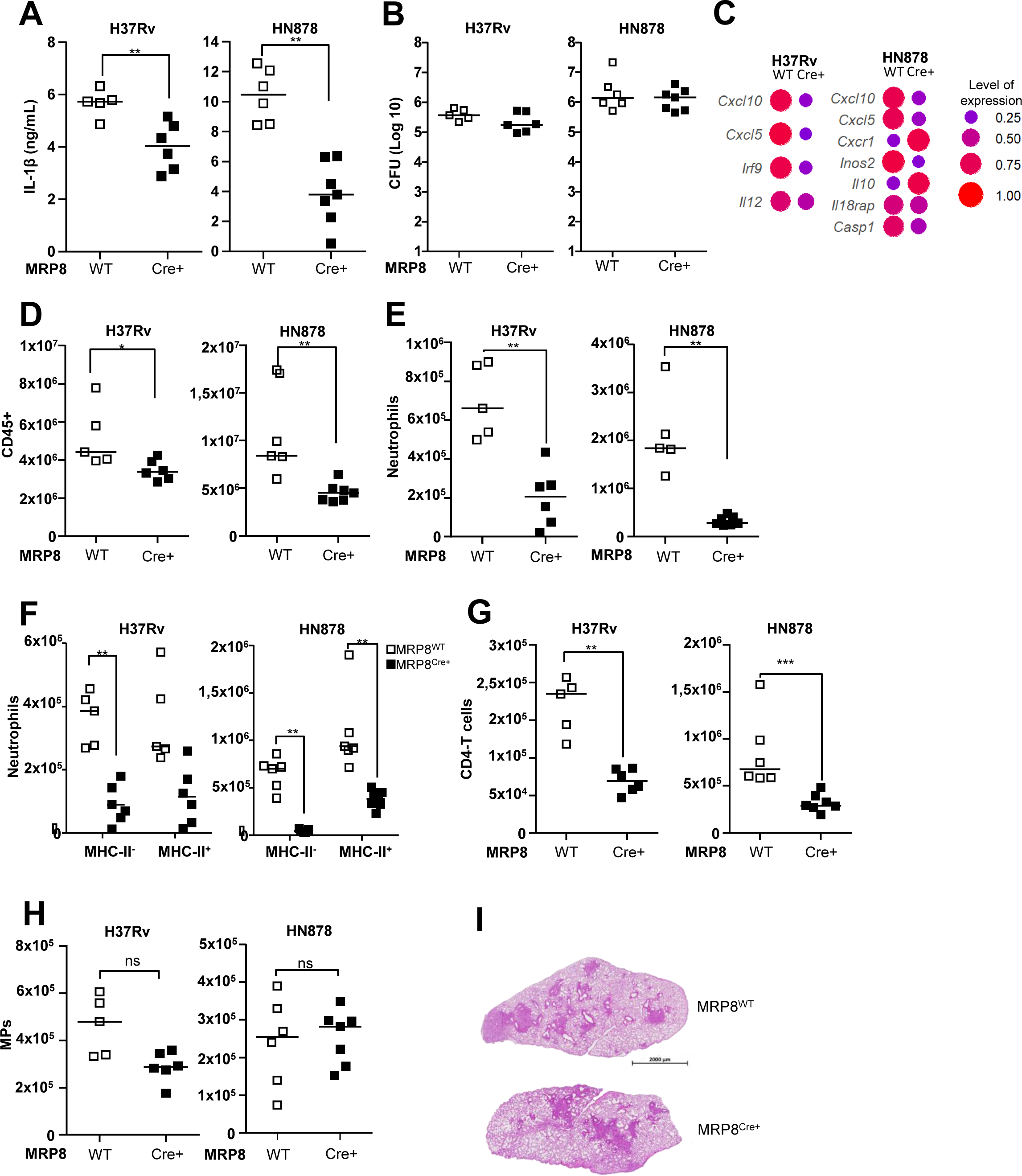
Caspase-dependent production of IL-1β by inflammatory neutrophils sustains lung inflammation. **(A-I)** *MRP8^Cre+^Csp1^flox^* or *MRP8^WT^Csp1^flox^*mice were infected with H37Rv or HN878 and the lungs harvested at 21 days. **(A)** IL-1b production quantified by ELISA in lung tissue homogenates. **(B)** Number (Log_10_) of CFUs for each animal. **(C)** Differential gene expression of a panel of 48 genes between *MRP8^Cre+^Csp1^flox^* and *MRP8^WT^Csp1^flox^* mice infected with H37Rv or HN878. The dot plot represents the normalized expression of significantly deregulated genes expressed as a normalized rate to compare the two groups. **(D-H)** Cells were analyzed by flow cytometry and the number of (**D**) CD45^+^ total leukocytes, **(E)** Ly6G^+^ total neutrophils, **(F)** MHC-II^-^ and MHC-II^+^ neutrophil subsets, **(G)** CD4^+^ T-cells, and **(H)** CD11c^+^ or ^-^ CD11b^+^ MPs were compared between *MRP8^Cre+^Csp1^flox^* and *MRP8^WT^Csp1^flox^*mice. **(I)** Representative section of hematoxylin/eosin lung staining for each group. Data are presented as individual data points and medians. (A-I) n = 5-7 mice per group; (A, B, D-I) two independent experiments; (C) one experiment. ∗*p* < 0.05, ∗∗*p* < 0.01, and ∗∗∗*p* < 0.001 by the Mann-Whitney test (A-E, G-H) and two-way ANOVA with a Bonferroni post-test (F).

After BCG instillation, total leukocyte numbers in the lungs were not significantly different between *MRP8^Cre+^Csp1^flox^* and *MRP8^WT^Csp1^flox^* mice (Fig. S3D), which correlated with no difference in IL-1β production. In response to H37Rv, total lung leukocyte numbers were 24% lower for *MRP8^Cre+^Csp1^flox^*than the *MRP8^WT^Csp1^flox^* controls (Fig 5D). In response to HN878 the decrease was 46%. This result confirms the direct role of neutrophilic Nlrp3 inflammasome activation in lung inflammation. Among leukocytes, 3.2 times fewer neutrophils (Fig. 5E) were recruited to the lungs of *MRP8^Cre+^Csp1^flox^*than *MRP8^WT^Csp1^flox^* mice in response to H37Rv and 6.3 times fewer were recruited in response to HN878. In H37Rv-infected mice, we observed five-fold fewer inflammatory neutrophils in the lungs of *MRP8^Cre+^Csp1^flox^*mice than the *MRP8^WT^Csp1^flox^* controls and approximately twofold - but not statistically significant-fewer regulatory neutrophils (Fig. 5F). On the contrary, in HN878-infected mice, the number of inflammatory neutrophils was 16 times lower in *MRP8^Cre+^Csp1^flox^* than *MRP8^WT^Csp1^flox^*mice, whereas the number of regulatory neutrophils was only 2.3 times lower (Fig. 5F). Thus, the absence of neutrophilic inflammasome activation had a greater impact on inflammatory neutrophils than regulatory neutrophils, which was even more marked in response to HN878 infection. Again, we observed much lower expression of PD-L1 on the surface of regulatory neutrophils in response to HN878 (MFI 2256) than H37Rv infection (MFI 6158) (Fig. S3E). However, the levels were similar in *MRP8^Cre+^Csp1^flox^* and *MRP8^WT^Csp1^flox^*mice, indicating that the ability of inflammatory neutrophils to produce IL-1β did not have an impact on PD-L1 expression of regulatory neutrophils. CD4 T-cell numbers were 2.7 times lower for *MRP8^Cre+^Csp1^flox^*than *MRP8^WT^Csp1^flox^* mice in response to H37Rv and 2.3 times lower in response to HN878 (Fig. 5G). There was no statistically significant difference in the recruitment of lung interstitial and alveolar MPs between *MRP8^Cre+^Csp1^flox^* and *MRP8^WT^Csp1^flox^*mice in response to H37Rv or HN878 (Fig. 5H). Despite the greater impact on cell recruitment on day 21 after infection with HN878, we did not observe differences in the surface area of the lung occupied by lesions between *MRP8^Cre+^Csp1^flox^* and *MRP8^WT^Csp1^flox^*mice (Fig. 5I), indicating that other regulatory circuits control lesions.

### Extremely susceptible IFN-**γ**R^-/-^ mice show dysregulation of both neutrophil subsets

Mendelian inherited susceptibility to mycobacteria involves IFN-γR and its signaling cascade (*34*). IFNγ-R^-/-^ mice are extremely susceptible to Mtb infection and this is linked to strong recruitment and dysregulated cell death of neutrophils (*25*). We infected IFN-γR^-/-^ mice with virulent H37Rv or avirulent BCG. We did not infect these extremely susceptible animals with hypervirulent HN878 for ethical reasons. As we did not observe any clinical condition in IFN-γR^-/-^ mice infected with BCG (data not shown), we did not pursue neutrophil analysis in these animals. Three weeks after infection with H37Rv, we observed macroscopic lesions in the lungs and livers of IFN-γR^-/-^ mice that were not seen in their WT counterparts (data not shown). The lungs of IFN-γR^-/-^ mice showed 2.3 times more CFUs than those of WT mice (Fig. 6A). In accordance with the high number of macroscopic lesions, histological analysis of the lungs of IFN-γR^-/-^ infected mice showed extensive, disorganized inflammatory cell infiltrates (Fig. 6B). The total surface occupied by lesions was 2.6 times higher for the IFN-γR^-/-^ than WT mice (Fig. 6C). As expected, (*25*) we observed twofold greater recruitment of total leukocytes to the lungs of IFN-γR^-/-^ than WT mice (Fig. 6D). This difference was mainly due to total neutrophils, which were 6.5 times more abundant in IFNγ-R^-/-^ than WT mice. The number of CD4^+^ T cells was also twofold higher in the lungs of IFNγ-R^-/-^ mice, whereas there was no difference in the number of CD8^+^ T cells (Fig. S4). Of note, inflammatory neutrophils represented 70% and regulatory neutrophils 30% of the total neutrophil influx in IFN-γR^-/-^ mice (Fig. 6D), whereas the neutrophil influx in WT controls was balanced between the inflammatory (41%) and regulatory (59%) subsets. The threefold higher level of IL-1β detected in the lungs of IFN-γR^-/-^ than WT mice (Fig. 6E) is consistent with the higher influx of inflammatory neutrophils. Higher inflammation was also indicated by the presence of 2.6 times more TNF (Fig. 6F) and 1.7 times more IL-6 (Fig. 6G) in the lungs of IFN-γR^-/-^ than WT mice. Lung tissue from the two mouse strains showed highly different transcriptional profiles in response to H37Rv infection (Fig. 6H). Genes such as *Cxcl1*, *Cxcr1*, *Cxcr2*, *Mmp7*, *Mmp8*, *Mmp9*, *Mpo*, and *S100a8*, were more highly expressed in IFN-γR^-/-^ than WT mice (Fig. 6I). Many genes, such as *Ilr1* and I*lr2*, which are highly expressed during inflammation, including by the neutrophils themselves, were also more highly expressed in IFN-γR^-/-^ than WT mice. By contrast, the expression of type I IFN-related genes was higher in WT than IFN-γR^-/-^ mice.

**Fig. 6.**
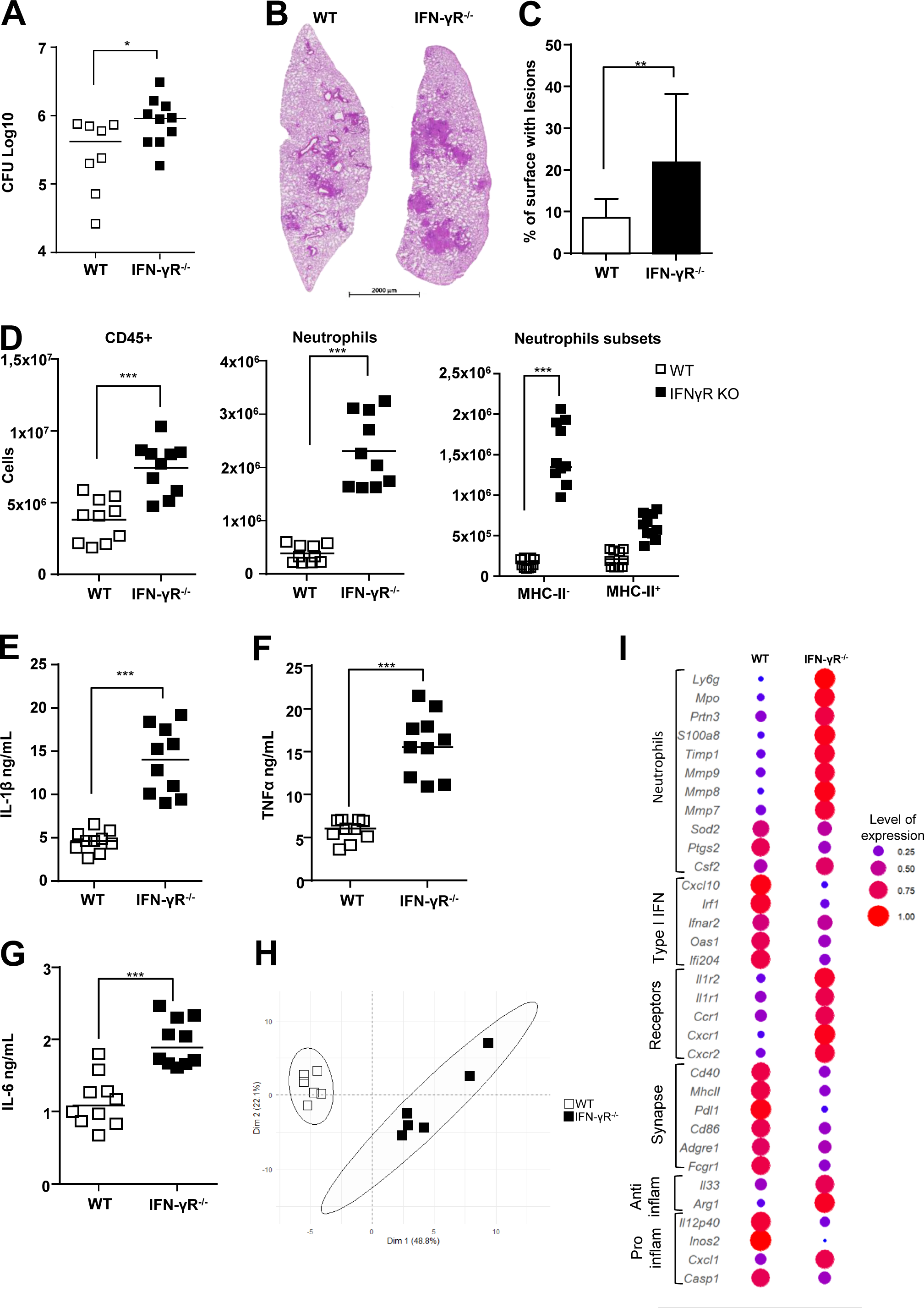
Extremely susceptible IFN-γR^-/-^ mice display dysregulation of both neutrophil subsets. C56BL/6 WT or IFN-γR^-/-^ mice were infected with H37Rv and the lungs harvested on day 21 for analysis. **(A)** Number (Log_10_) of CFUs for each animal. **(B)** Representative section of hematoxylin/eosin lung staining for each group and **(C)** mean percentage of lung surface occupied by lesions. **(D)** Cells were analyzed by flow cytometry and the number of CD45^+^ total leukocytes, Ly6G^+^ total neutrophils, and MHC-II^-^ and MHC-II^+^ neutrophil subsets were compared between the two groups of mice. **(E-G)** Cytokine production analyzed by ELISA in lung tissue homogenates. **(H-I)** Gene expression of a panel of 48 genes in the lungs assessed by Fluidigm Biomark. **(H)** mRNA expression was normalized to the expression of three housekeeping genes and to the uninfected group to calculate the ΔΔCt. Principal component analysis (PCA) was performed on the ΔΔCt values. The two first dimensions of the PCA plot are depicted. (**I**) The dot plot represents the normalized expression of significantly deregulated genes expressed as a normalized rate to compare the C56BL/6 WT and IFN-γR^-/-^ mice. (A-I) Data are presented as the mean of n = 5-7 mice per group; (A-G) pooled data from two independent experiments; (G-H) analysis of one experiment. Graphs are presented as individual data points and medians (C, median with range). (G) Data are presented as the mean of n = 6 mice per group. ∗*p* < 0.05, ∗∗*p* < 0.01, and ∗∗∗*p* < 0.001 by the Mann-Whitney test (A-I) and two-way ANOVA with a Bonferroni post-test (D, neutrophil subsets).

### Hyperinflammation in IFN-**γ**R^-/-^ mice is relieved by IFN-**γ**R^+^ regulatory neutrophils

The genes for which the expression was higher in the lungs of WT than those of IFN-γR^-/-^mice in response to H37Rv infection included *Mhc-II*, *CD274*, *Cd86*, and *Cd40* (Fig. 6I), which are all involved in the synapse between antigen-presenting cells and T cells. As these mice show a hyperinflammatory profile, we investigated the impact of the IFN-γR on regulatory neutrophils. The level of MHC-II expression was not affected by the absence of the IFN-γR (Fig. 7A). However regulatory neutrophils from IFN-γR^-/-^ mice lost PD-L1 surface expression, showing levels similar to those of inflammatory MHC-II^-^ neutrophils from WT animals (Fig. 7B). Moreover, the proportion of MHC-II^+^ neutrophils that expressed low levels of PD-L1 in IFN-γR^-/-^ mice dropped to 30%, whereas 90% of MHC-II^+^ neutrophils highly expressed PD-L1 in WT animals (Fig. 7C). We enriched for lung regulatory neutrophils from IFN-γR^-/-^ or WT mice 21 days after H37Rv infection by magnetic sorting. Strikingly, IFN-γ-R^-/-^ regulatory neutrophils completely lost the ability to suppress OT-II-cell proliferation (Fig. 7D) and IFN-γ production *ex-vivo* (Fig. 7E), showing that the control exerted by regulatory neutrophils on T cells is dependent on the IFN-γR. Thus, we hypothesized that lethal inflammation in Mtb-infected IFN-γR^-/-^ mice was linked to strong recruitment of inflammatory neutrophils and less efficient control of inflammation by regulatory neutrophils due to PD-L1 downregulation. We tested this hypothesis by harvesting PD-L1^hi^ regulatory neutrophils from BCG-infected WT mice and transferring them into H37Rv-infected IFN-γR^-/-^ mice on day 18 post-infection. On day 21, we euthanized these mice, as well as the two control groups, H37Rv-infected WT and mock-treated IFN-γR^-/-^ mice (Fig. 7F), and assessed the TB physiopathology in the lungs. We did not observe any significative differences in CFU counts between the groups at this time point (Fig. S5A). However, transfer of PD-L1^hi^ regulatory neutrophils relieved inflammation in the lung tissue from IFN-γR^-/-^ mice (Fig. 7G), although the difference in total lung-surface occupied by lesions among mock-treated and regulatory neutrophil transferees did not reach statistical significance (Fig. 7H). Nonetheless, the dampening of inflammation was also indicated by a significant reduction in total leukocyte numbers (Fig. 7I), in particular, those of neutrophils (Fig. 7J) and T cells (Fig 7K), including both CD8^+^ (Fig. S5B) and CD4^+^ T cells (Fig. S5C). The transfer of regulatory neutrophils to IFN-γR^-/-^ infected mice dampened the strong recruitment of inflammatory neutrophils observed in mock-treated animals (Fig. 7L) and the PD-L1 MFI of MHC-II^+^ neutrophils increased significantly (Fig. 7M). Consistent with these results, we measured 1.4-fold less IL-1β production in the lung tissue from transferees than mock-treated animals (Fig. 7N). IFN-γ levels were also 1.8-fold lower (Fig. 7O) and those of TNF remained unchanged at this time point (Fig. S5D).

**Fig. 7.**
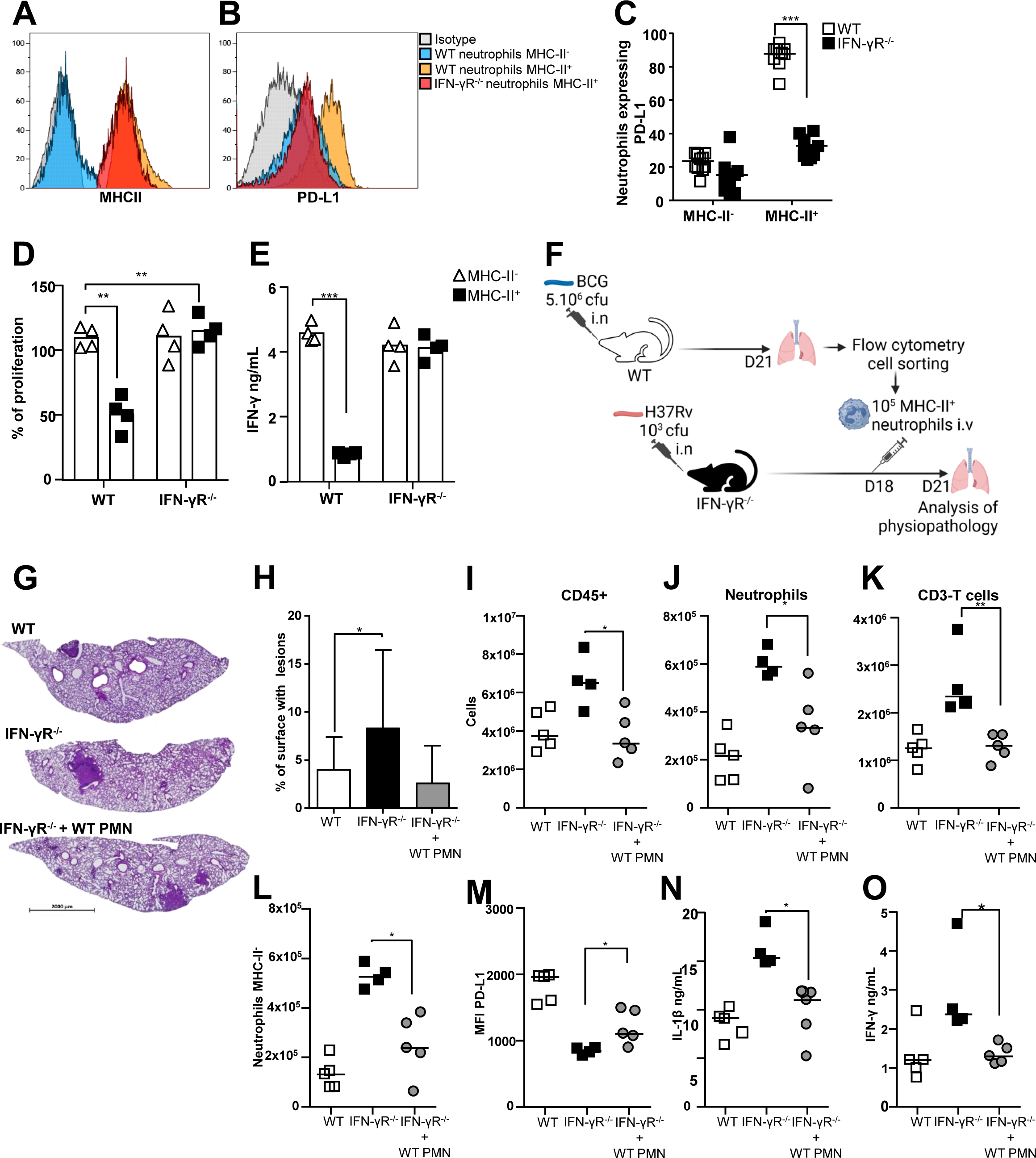
Hyperinflammation in IFN-γR^-/-^ mice is relieved by IFN-γR^+^ regulatory neutrophils. C56BL/6 WT or IFN-γR^-/-^ mice were infected with H37Rv and the lungs harvested on day 21 for analysis. Neutrophil subsets analyzed by flow cytometry for **(A)** MHC-II and **(B)** PD-L1 surface expression. **(C)** Percentage of neutrophils expressing PD-L1 on the surface among the two MHC-II^-^ and MHC-II^+^ neutrophil subsets in the two mouse groups. **(D-E)** On day 21, MHC-II^-^ and MHC-II^+^ neutrophil subsets were enriched by magnetic beads from the lungs of the two groups of mice and mixed with OT-II cells. **(D)** Percentage of OT-II splenocyte proliferation and **(E)** IFN-γ production in the presence of each neutrophil subset calculated based on the response of OT-II splenocytes to the Ova peptide only. **(F)** Schematic representation of the transfer of [MHC-II^+^, PD-L1^hi^] regulatory neutrophils purified from the lungs of BCG-infected WT mice into IFN-γR^-/-^ mice infected with H37Rv 18 days before. The three H37Rv-infected groups harvested on day 21 were WT control mice and IFN-γR^-/-^ mice that were mock-treated or to which WT regulatory neutrophils were transferred. **(G)** Representative section of hematoxylin/eosin lung staining for each group and **(H)** the percentage of lung surface occupied by lesions analyzed. **(I-O)** Cells were analyzed by flow cytometry to determine the number of **(I)** CD45^+^ total leukocytes, **(J)** Ly6G^+^ total neutrophils, **(K)** T-cells, and **(L)** MHC-II^-^ inflammatory neutrophils. **(M)** Comparison of the PD-L1 mean fluorescence intensity on MHC-II^+^ regulatory neutrophils between the three groups analyzed by flow cytometry. **(N)** IL-1β and **(O)** IFN-γ production measured in lung tissue homogenates by ELISA. (A-B) Histograms are representative of two independent experiments, n = 6 or 4; (C) pooled data from two independent experiments, n = 10; (D-E) pooled data from two independent experiments (n = 4); (G-O) data from one experiment, n = 4-5. Data are presented as individual data points and medians (H, with range). ∗*p* < 0.05, ∗∗*p* < 0.01, and ∗∗∗*p* < 0.001 by two-way ANOVA with a Bonferroni posttest (C-E) and the Mann-Whitney test (G-O).

## DISCUSSION

TB physiopathology in the lung is characterized by a delicate balance between pro- and anti-inflammatory mechanisms controlled both by the host and bacilli. Neutrophils play key roles in this balance. Here, we show that two distinct subsets are recruited to the lungs in response to mycobacterial infection in a mouse model. The inflammatory neutrophil subset produces caspase 1-dependent IL-1β and acts as an accelerator of local inflammation in response to virulent mycobacteria by maintaining a vicious circle of inflammatory neutrophils and CD4 T cells. The regulatory neutrophil subset is able to dampen inflammation by blocking T-cell proliferation and IFN-γ production. IFNγ-R-dependent expression of PD-L1 on regulatory neutrophils is critical for the braking function. Regulatory neutrophils are less affected than inflammatory neutrophils by the absence of neutrophil-derived IL-1β, suggesting differential regulation mechanisms. We propose (Fig. 8) that these two subsets are involved in a “brake/accelerator” inflammation circuit in the lungs during TB infection. Moreover, the two brake and accelerator pedals could represent a means for hypervirulent Mtb strains to manipulate the host’s immune system and establish a successful infection.

**Fig. 8.**
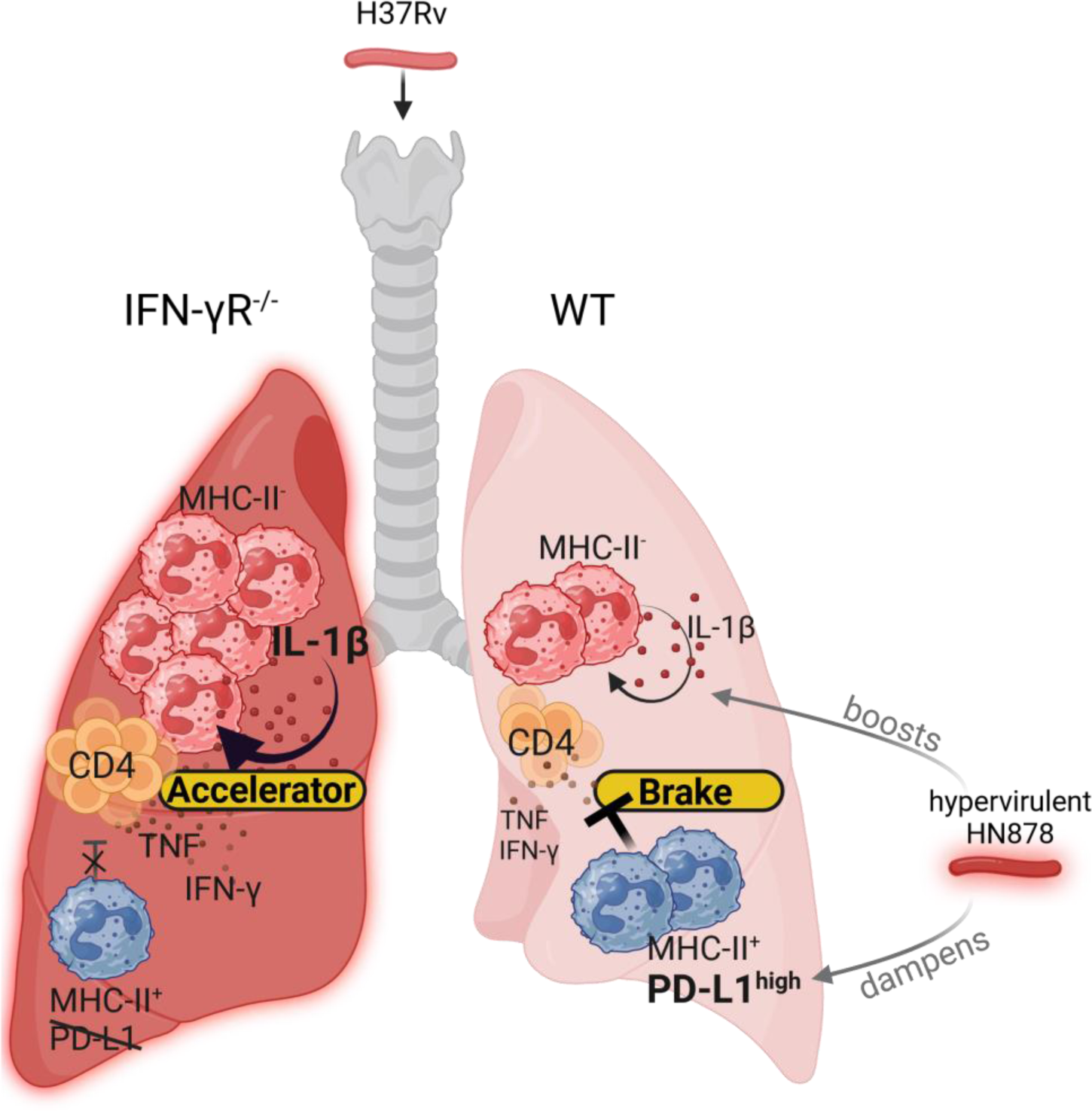
Graphical abstract of the dual “brake” and “accelerator’ functions exerted by the two neutrophil subsets on Mtb infection in the lung. Two functionally distinct neutrophil subsets are recruited to the lungs in response to mycobacterial infection. Classic [MHC-II^-^, PD-L1^lo^] neutrophils produce high levels of inflammasome-dependent IL-1β in the lungs in response to virulent mycobacteria and act as an “accelerator” of deleterious inflammation, which is highly exacerbated in IFN-γR^-/-^ mice. Conversely, regulatory [MHC-II^+^, PD-L1^hi^] neutrophils act as a “brake” of inflammation by suppressing T-cell proliferation and IFN-γ production. This regulation depends on PD-L1 surface expression, controlled by IFN-γR signaling in neutrophils. The hypervirulent HN878 strain from the Beijing genotype curbs PD-L1 expression by regulatory neutrophils, which abolishes the brake pedal and drives deleterious hyperinflammation in the lungs.

IL-1β is a double-edged sword during TB infection that must be tightly controlled. It is involved in strong neutrophil recruitment to the lungs during severe TB (*12, 35, 36*) and is a target for host-directed therapies (*37*). Neutrophils produce IL-1β during Mtb infections and we demonstrated here that caspase-1 dependent cleavage of pro-IL1β occurs in neutrophils, in addition to protease-dependent mechanisms (*38*). Avirulent BCG can trigger caspase-1-dependent IL-1β production by neutrophils *in vitro*, showing that the major virulence factor ESAT-6, which is present in Mtb and absent from BCG and required to trigger NLRP3 in MPs (*39*), is dispensable in neutrophils. However, *in vivo*, caspase-1 dependent IL-1β production by neutrophils was only induced by Mtb and not BCG, indicating that other regulatory pathways are involved in inflammasome activation in the lungs. Three weeks following Mtb infection of mice bearing caspase-1 defective neutrophils, we observed a 30% to 64% reduction of IL-1β levels in the lungs depending on the virulence of the strain and a coincident 3.2-to-6.3-fold reduction in total neutrophil recruitment, underlining the importance of the caspase-1-dependent pathway for the inflammatory loop involving neutrophils in the lung. In our study, restricted to one timepoint corresponding to early orchestration of the adaptive T-cell response in the lungs (*16, 40*), we observed that caspase-dependent IL-1β production mainly affected the recruitment of inflammatory neutrophils and CD4 T cells. The link between IFN-γ-producing CD4 T cells and excessive neutrophilia during clinical manifestations of TB has been clearly established (*41*) and we observed unrestricted recruitment of IL-1β-producing inflammatory neutrophils in IFN-γR^-/-^ mice that correlated with elevated levels of CD4 T cells and IFN-γ and highly lesioned lungs. Thus, IL-1β-producing inflammatory neutrophils are more involved than regulatory neutrophils in severe forms of TB. Although we did not observe a major impact of caspase-dependent IL-1β production by inflammatory neutrophils on control of the bacilli or lesion formation at the early timepoint of our studies, we believe that other timepoints should be examined in *MRP8^Cre+^Csp1^flox^* mice to gain a better understanding of caspase-1-versus protease-dependent mechanisms of IL-1β production by neutrophils.

In highly susceptible IFN-γR^-/-^ mice, we observed that the strong neutrophilia was driven by two conjugated paths, dysregulated recruitment of IL-1β-producing inflammatory neutrophils and dysfunction of PD-L1^hi^ regulatory neutrophils, which could be alleviated by the transfer of competent WT regulatory neutrophils. The immune checkpoint inhibitor PD-L1 was critical for the function of regulatory neutrophils, akin to neutrophils present in cancer (*42*), which foster immune suppression in hepatocellular carcinoma (*43, 44*) and gastric cancer (*45*). Competent PD-L1^hi^ regulatory neutrophils were also recruited to the lungs in response to avirulent BCG-infection, indicating that the acquisition of this function did not fully depend on mycobacterial virulence. Recently, PD-L1^+^ neutrophils were described in two acute disorders, sepsis (*46, 47*) and cutaneous burn injury (*48*), as well as during chronic infections in cutaneous (*49*) or visceral (*50*) leishmaniasis. During *Candida albicans* infection, PD-L1^+^ neutrophils decrease antifungal immunity by retaining the pool of microbiocidal neutrophils in the bone marrow (*51*). We found that the IFN-γR was required for PD-L1 expression and suppression of CD4 T cells. Similarly, human (*52*) and mouse neutrophils need to be exposed to IFN-γ to express PD-L1 and suppress T-cells during endotoxemia (*46*). Moreover, we found that the transfer of WT regulatory neutrophils that expressed PD-L1 into Mtb-infected IFN-γR^-/-^ mice alleviated exuberant lung neutrophilia and lesions in these extremely susceptible animals.

The tremendous success of Mtb as a pathogen can be explained by its co-evolution with that of the host. Strains from the Beijing family are among the most successful, as demonstrated by their global distribution and the recurrent outbreaks they cause (*53*). This success is partially due to their exquisite ability to manipulate the host’s immune system. The peculiar cell-wall composition allows the Beijing strains to immunosuppress the innate immune response (*54*), especially in microaerophilic or anaerobic environments (*55*), such as that encountered in the granuloma. Here, we confirm the hypervirulence of Beijing prototype strain HN878 in C57BL/6 mice, with neutrophil-driven lung inflammation (*29, 30*). A neutrophil-driven type I IFN response has been shown to lead to a poor prognosis for TB patients (*56*) and mice (*57*). Our data, restricted to one timepoint in C57BL/6 mice, indicate better induction of the type I IFN pathway in the lungs by the less virulent H37Rv strain than hypervirulent HN878. However, unlike H37Rv, HN878 was able to fuel the neutrophil influx towards recruitment of the regulatory subset, with diminished PD-L1 expression. Mtb Beijing strains induce regulatory T-cell expansion (*58, 59*) better than lab-adapted strains. They also favor recruitment of myeloid-derived suppressor cells producing IL-10, which could limit excessive lung damage (*29*). Of note, we also observed higher expression of *Il10* and *Arg1* in the lungs of mice infected with HN878 than those infected with H37Rv. Our most striking finding was the ability of HN878 to recruit a neutrophil compartment biased towards regulatory neutrophils, which expressed three-to five-fold less PD-L1 on their surface than less virulent lab-adapted strains. We observed a similar difference at the transcriptional level in lung tissue. As PD-L1 is widely expressed both by myeloid and non-hematopoietic cells (*60*), it is possible that control of this important immune checkpoint by HN878 occurs at several levels at the site of infection. Further studies are required to better dissect the mechanisms used by diverse Mtb strains to finely tune PD-L1 expression and how they relate to the functional consequences of infection. We believe that regulatory neutrophils acting as a “brake pedal” represent yet another weapon in the arsenal of Beijing strains to manipulate the immune system and establish successful infection.

Our study had two principal limitations. First, the function of regulatory neutrophils was not assessed in TB patients. Although PD-L1^+^ neutrophils have been found in TB patients (*61*), this subset is yet to be investigated in the lungs of humans and non-human primates. Second, although the C57BL/6 mouse model is the most widely used because it allows mechanistic studies in genetically modified mice, granulomas are not well-formed in the lungs in response to Mtb infection in this model. Therefore, our next step will be to examine the contribution of the two neutrophil subsets to granuloma formation in C3HeB/FeJ mice (*18*).

In conclusion, our results add a new layer of complexity to the multiple functions exerted by different neutrophil subsets during TB and emphasize their key role as partners of the immune response. Inflammatory neutrophils are certainly “foes”, worsening TB pathogenesis in the lung. It is yet to be determined whether regulatory neutrophils are “friends” and associated with a good prognosis for TB patients. Recent demonstration of the importance of the PD-1/PD-L1 axis in the control of TB (*6*) and the data we report here on regulatory PD-L1^hi^ neutrophils open new avenues to explore the role of this subset in the granuloma microenvironment in humans. As neutrophils, in general, are an important target for the development of new host-directed therapies (*62, 63*), it is urgent to reconsider the complexity of these cells to better target pharmaceutical and immune interventions in TB.

## MATERIALS AND METHODS

### Experimental design and justification of the sample size

Mice were bred at the specific pathogen-free animal (SPF) facility Plateforme Infectiologie Experimentale (PFIE, U1277, INRAE, Center Val de Loire). One week before *in vivo* experiments, mice were moved from the SPF breeding area to the ABSL3 area to acclimate them. For infections with Mtb, mice were placed in biological safety cabinets. Mice were housed in groups of 4 to 5 per cage and randomly distributed. *MRP8^Cre+^Csp1^flox^*and *MRP8^WT^Csp1^flox^* mice were littermates. Groups always contained at least four individuals to allow statistical assessment of the data using non-parametric Mann Whitney tests. Results were not blinded for analysis excepted for the RNAseq analysis. The number of biological replicates and experiments are indicated in the figure legends.

### Ethics statement

Experimental protocols complied with French law (Décret: 2001–464 29/05/01) and European directive 2010/63/UE for the care and use of laboratory animals and were carried out under Authorization for Experimentation on Laboratory Animals Number D-37-175-3 (Animal facility UE-PFIE, INRAE Centre Val de Loire). Animal protocols were approved both by the “Val de Loire” Ethics Committee for Animal Experimentation and the French Minister of Higher Education, Research and Innovation. They were registered with the French National Committee for Animal Experimentation under N° APAFIS #35838-2022031011022458.v5.

### Mice

*MRP8^Cre+^Csp1^flox^* mice have been previously described (*26*). Because introduction of the *cre* gene encoding the recombinase under control of the MRP8 promoter in both alleles of the C57BL/6 mouse chromosome was lethal, we bred and screened mice to obtain *MRP8^Cre+^Csp1^flox^*mice in which one allele carried the CRE recombinase while the other did not. In these animals, expression of the recombinase under the MRP8 promoter induced excision of the *Csp1*-encoding genes in 100% of the neutrophils. Control *MRP8^WT^Csp1^flox^* mice did not carry the recombinase under the control of the MRP8 promoter and neutrophils were able to cleave pro IL-1β. All mice were bred in-house, except OT-II mice, which were purchased from Janvier Biolabs.

### Bacterial strains and growth conditions

All mycobacterial strains *(M. bovis* BCG strain WT 1173P2 Pasteur, Mtb strains H37Rv and HN878) were cultivated for 12 days in Middlebrook 7H9 broth (Becton Dickinson, USA) supplemented with 10% BBL™ Middlebrook ADC enrichment (BD, USA) and 0.05% Tween 80 (Sigma-Aldrich, USA), aliquoted, and frozen at -80°C in 7H9 medium containing 10% glycerol. Bacterial suspensions for infection were prepared in PBS from quantified glycerol stock solutions. To enumerate the number of CFUs from the middle right lung lobe, tissue was homogenized, and serial dilutions were plated on supplemented 7H11 plates as previously described (*16*).

### Intranasal infection and treatments

Mice anesthetized by i.p. injection of a ketamine/xylazine cocktail received 5×10^6^ CFUs of BCG Pasteur or 10^3^ CFUs of Mtb H37Rv or HN878 in 20 μL in each nostril. For total neutrophil depletion experiments, C57BL/6 mice received 200 μg anti-Ly6G antibody (clone 1A8, Biolegend) via the i.p. route on days 15, 17, and 19 following BCG inoculation. Control mice were injected with the same quantity of IgG2b Ab (Biolegend). For neutrophil transfer experiments, MHC-II^+^ neutrophils were isolated from the lungs of BCG-infected mice harvested on day 21. IFN-γR^-/-^ and control C57BL/6 mice that were infected with H37Rv 18 days before were injected i.v. with 1.5×10^5^ MHC-II^+^ neutrophils. Mice were euthanized on day 21 for analysis of the lungs. All mice were euthanized by pentobarbital administration at the time post-infection indicated in the figure legends.

### Preparation of neutrophils and macrophages from bone marrow

Femurs were harvested from six-week-old mice (WT, *MRP8^WT^Csp1^flox^*and *MRP8^Cre+^Csp1^flox^*) bred at the PFIE animal facility. Femurs from *Aim2*^-/-^, *Gsdmd*^-/-^, *Nlrp*3^-/-^, and *Csp1/11*^-/-^ mice were kindly donated by Valérie Quesniaux (INEM UMR7355 CNRS, University of Orleans, France) and the of *Csp1*^-/-^ mice by Sergio Costa (Universidade Federal de Minas Gerais, Belo Horizonte, Brazil). Neutrophils were directly purified from bone marrow by anti-Ly-6G magnetic positive selection (Miltenyi Biotec), as previously described (*64*). Neutrophils of > 95% purity were obtained as assessed by microscopy after May-Grünwald-Giemsa staining. Viability by trypan blue exclusion was 98%. MPs were obtained after culture with 30% L929 cell-conditioned medium as a source of macrophage colony-stimulating factor. Cells used on day 10 for infectivity and cytokine assays were suspended in complete medium without antibiotics, as previously described (*64*). Macrophages (1×10^5^/well) or neutrophils (1×10^6^/well) were plated in P96 plates and infected overnight with BCG at an MOI of 10 or Mtb at an MOI of 1 or stimulated overnight with 100 ng LPS (from *E. Coli* 011: B4, Sigma) and 10 µM nigericin sodium salt (Sigma) added 1 h before harvesting the cells and supernatants.

### Lung cell preparation and flow cytometry

Briefly, euthanized mice were perfused with PBS and the left lung lobes digested for 1 h with collagenase D (1.5 mg/ml, Roche) and DNAse A (40 U/ml, Roche) before filtering cells through a 100 μM nylon cell strainer (BD Falcon). For extracellular staining, cells were incubated 20 min with 2% total mouse serum and labeled in PBS supplemented with 5% FCS and 0.1% total mouse serum with antibodies against the surface markers, all from BD Biosciences (listed in Supplementary Table 1). Intracellular mature IL-1β production was measured using anti IL-1β biotin conjugated antibody (Rockland) after treatment for 2 h at 37°C with 5 μg/ml brefeldin A (Sigma-Aldrich). Cells were washed and fixed with BD cell fix diluted 4X in PBS. Data were acquired on a LSR Fortessa™ X-20 Flow cytometer (Becton Dickinson) and the results analyzed using Kaluza software (Beckman Coulter).

Lung regulatory or inflammatory neutrophils were prepared from the lungs of C57BL/6 mice on day 21 post-infection with Mtb or BCG. For Mtb, lungs were digested and the neutrophils isolated by magnetic bead selection using the untouched neutrophil isolation kit according to the manufacturer’s instructions (Miltenyi). MHC-II-positive magnetic bead selection was performed on the unlabeled neutrophil-rich fraction. MHC-II^+^ regulatory neutrophils were separated from MHC-II^-^ inflammatory-neutrophils using anti-MHC-II PE conjugated Ab (BD Biosciences) and anti-PE beads (Miltenyi). Viability by trypan blue exclusion was 95%. For BCG, total neutrophils [CD11b^+^, Ly-6C^+^, Ly-6G^+^], classic neutrophils [CD11b^+^, Ly-6C^+^, Ly-6G^+^, MHCII^-^], or regulatory neutrophils [CD11b^+^, Ly-6C^+^, Ly-6G^+^, MHC-II^+^] were sorted on a MoFlo AstriosEQ high speed cell sorter (Beckman Coulter) as previously described (*19*). Neutrophil subsets of > 99% purity were obtained in each fraction. Neutrophils were recovered in complete medium and immediately processed for single-cell RNAseq analysis (total neutrophils) or transcriptomic analysis or neutrophil transfer (neutrophil subsets).

### Measurement of T-cell suppressive activity of neutrophils

The T-cell suppressive activity of neutrophils was measured as previously published (*19*). Briefly, total splenocytes from OT-II mice were collected, homogenized to single-cell suspensions through nylon screens, and resuspended in RPMI medium (Gibco) supplemented with 10% decomplemented fetal bovine serum (Gibco), 2 mM L-glutamine (Gibco), 100 U penicillin, and 100 μg/ml streptomycin (Gibco). Then 10^5^ cells/well were distributed in a 96-well round bottom plate (BD Falcon). OT-II splenocyte proliferation was induced by the addition of 2 μg/ml of the OVA peptide 323-339 (PolyPeptide Group). Purified neutrophils were added to the cultured splenocytes at a ratio of 1:10 in a final volume of 200 µl. Wells without neutrophils were used as a reference for maximal proliferation. Cell proliferation was quantified after three days of culture using CyQUANT Cell Proliferation Assay tests (Thermo Fisher) according to the manufacturer’s instructions. The role of PD-L1 in the suppression mechanism was assessed by incubating sorted neutrophils for 1 h with 50 µg/ml anti-PD-L1 Ab (Tecentriq®, atezolizumab) or a human IgG1 isotype control before mixing with OT-II splenocytes. Cell proliferation was quantified after three days of culture using CyQUANT Cell Proliferation Assay tests (Thermo Fisher) according to the manufacturer’s instructions.

### Medium-throughput and single-cell RNA sequencing of neutrophils

For medium-throughput analysis of gene expression in neutrophils, total RNA was extracted from FACS-sorted neutrophils from the right accessory lung lobe homogenized using Lysing matrix D tubes from MP Biomedicals and a Precellys® using a NucleoSpin RNA kit with DNase treatment (Macherey Nagel). Total RNA was reverse transcribed using an iScript™ Reverse Transcriptase mix (Biorad) and gene expression assessed using a BioMark HD (Fluidigm) according to the manufacturer’s instructions or a LightCycler® 480 Real-Time PCR System (Roche). The annealing temperature was 62°C. All primers are listed in Supplementary Table 2. Data were analyzed using Fluidigm RealTime PCR software or Lc480 software to determine the cycle threshold (Ct) values. Messenger RNA (mRNA) expression was normalized against the mean expression of three housekeeping genes for each sample to obtain the ΔCt value. Infected samples were normalized against uninfected samples (ΔΔCt). Relative gene expression was calculated according to the formula RQ = 2^-ΔΔCt^. Dot plots were created using Rstudio for differentially expressed genes between two groups (Mann Whitney) by normalizing the fold change of each group to the total fold expression for each gene [normalized-rate = fold change group 1 / (foldchange group 1 + foldchange group 2]. This normalized rate is represented as spot plots when the transcriptomes of two groups are compared.

For single-cell analysis, viable total Ly6G**^+^** neutrophils were sorted using a MoFlowAstrios high-speed cell sorter. Within 1 h after sorting, cells were encapsulated with barcoded Single Cell 3L v3.1 Gel Beads and a Master Mix to form a Gel Beads-in-emulsion using the 10X Genomics Chromium technology. Approximately 12,000 cells were used. The Single Cell 3’ libraries were then generated as recommended by the manufacturer (10x Genomics). The libraries were equimolarly pooled and sequenced (paired-end sequencing) using one lane of an Illumina NovaSeq6000 device (IntegraGen, France), yielding a total of 640 million reads. Raw sequencing data are available under the following BioProject accession number PRJNA1026083. Fastq files were analyzed the sequences and aligned against those of the *Mus musculus* genome mm10 (GRCm38-release 98) using the cellrangercount pipeline of CellRanger software (v6.0.2). Downstream analyses were performed using R (v4.3.0), Rstudio and the following packages: Seurat (v4.4.0), SingleR (v2.2.0), celldex(v1.10.1). Quality controls first included empty droplets and doublets removal. Then, only droplets with at least 100 features and 1000 counts were retained. Normalization was done using the LogNormalize method and the 3000 most variable features.

Dimension reduction method was performed by Principal Component Analysis (PCA) retaining the first 9 principal components with a resolution of 0.8 to identify the clusters with FindClusters.

Cell-type inference was performed using SingleR and the Immunologic Genome Project database retrieved through celldex package.

## ELISA

Cell culture supernatants or right caudal lung lobe tissue, homogenized as above and supplemented with anti-proteases (ROCHE), were passed through 0.2 µm filters and either processed immediately or frozen at -20°C. Cytokine levels were measured by ELISA using kits (R&D Systems) according to the manufacturer’s instructions. Absorbance was measured on a Multiskan C plate reader (ThermoFisher).

### Histology

The right cranial lung lobe was fixed in 4% paraformaldehyde for 48 h. Subsequently, the tissue was dehydrated and stored in 70% ethanol before being embedded in paraffin. Five micrometer sections were cut and stained with hematoxylin and eosin (H&E) using a slide stainer (ST5020; Leica Biosystems, Nußloch, Germany). All slides were scanned on a slide scanner (AxioScan Z1; Zeiss, Oberkochen, Germany). Morphological analyses were performed using QuPath software (*65*); available at https://qupath.github.io/, version 0.4. Briefly, airway lesions were quantified using a semi-automated macro. The total area of tissue was automatically measured using a threshold and the lesions blindly measured manually for all slides. Data for each mouse consist of the mean of eight sections, cut every 100 µm, to accurately represent the whole lung.

### Western blots

Neutrophils (5×10^6^) s were seeded in six-well plates in Opti-MEM Glutamax medium (Gibco) at 37°C. Cells were infected for 5 h with BCG at a MOI of 10 or 20 or stimulated with 500 ng LPS from *E. coli* 011: B4 (Sigma) and 10 µM nigericin sodium salt (Sigma) added 45 min before the end of the incubation. Then, 1.5 mM AEBSF anti-protease (Sigma) was added to the wells and the supernatants clarified by centrifugation at 1500 x g, 10 min. Neutrophil lysates were prepared as previously described (*66*). For western blotting, whole cell lysates and supernatants were heated for 5 min at 95°C with 4X Laemmli buffer (Biorad) and the samples loaded on a 12% SDS-PAGE gel before transfer onto a nitrocellulose membrane using a TransBlot Turbo System (Biorad). After saturation in 5% non-fat milk/ TBS-0.1%Tween, membranes were incubated overnight at 4°C with the primary antibodies listed in Supplementary Table 2. After washing, the membranes were incubated 1 h at room temperature with secondary antibodies. Bands were visualized using Clarity Max ECL (Biorad) on a Fusion FX imaging system (Vilber Lourmat). Total proteins were measured after stripping for 12 min with Restore WB Stripping Buffer (Thermo), using GAPDH (D16H11) XP rabbit mAb (Cell Signaling Technology).

### Statistical analysis

Individual data and medians are presented in the Figures. Statistical analyses were performed using Prism 6.0 software (GraphPad). Analyses were performed on data from 2 to 6 independent experiments. Mann Whitney non-parametric tests or two-way ANOVA tests were used. For figure 6H, principal component analysis was performed using R studio with the factoMineR package. For figure 7H, the Rcmd plugin was used to analyze data as a stratified test. A paired non-parametric two tailed K-Sample Fisher-Pitman Permutation test was used to analyze data, with a Monte Carlo resampling approximation. Represented p-values are: ∗*p* < 0.05, ∗∗*p* < 0.01, and ∗∗∗*p* < 0.001.

## Acknowledgments

The mouse team of the PFIE (INRAE, Nouzilly), especially Corinne Beaugé, Jérôme Pottier, Emilie Lortscher and Laetitia Mérat, is warmly acknowledged for attentive mouse care and prompt reply to all our requests for mouse experiments. We thank Valérie Quesniaux (INEM, UMR7355 CNRS, Université d’Orléans, France) who generously provided bone marrow from *Aim2^-/-^, Gsdmd^-/-^, Nlrp3^-/-^, Csp1/11^-/-^* mice, and for helpful discussions. Alix Sausset and Christelle Rossignol, from the IMI team (ISP, INRAE, Nouzilly) are acknowledged for their assistance respectively with neutrophil cell sorting and histology. Warm thanks to Sonia Lacroix-Lamandé and the AIM team (ISP, INRAE, Nouzilly) for primers for the Fluidigm Biomark. We thank Roland Brosch (Institut Pasteur, Paris) who kindly provided the HN878 Mtb strain, and Sébastien Leclercq (ISP, INRAE, Nouzilly, France) who verified its genome sequence. Anti-PD-L1 Ab (Tecentriq®, atezolizumab) was graciously given by André Rieutord and Hail Aboudagga (Gustave Roussy Cancer Campus, Villejuif, France). Finally, we deeply thank Mustapha Si-Tahar and Hervé Watier (CEPR, UMR 1100, INSERM, Université de Tours, France) for helpful discussions.

## Author contributions

Conceptualization: NW, EDD, AR, BB, and CP

Methodology: EDD, BB, FK, and JP

Software: FK

Validation, BB, FC, and JSF

Formal Analysis: EDD, BB, AR, PG, JP, and FK

Investigation: EDD, BB, FC, JSF, and JP

Visualization: NW, EDD, BB, AR, FC, and JSF

Writing-original draft: NW, EDD, BB, and AR

Writing-review and editing: NW, EDD, BB, AR, SCO, PG, and CP

Resources: ML and SCO

Funding acquisition: NW

Project administration: NW, EDD, and AR

Supervision: NW, EDD, and SCO

## Data and material availability

All data are available in the main text or supplementary materials. Transcriptomic data are available using the BioProject accession number PRJNA1026083.

