## Supplementary material for "Neutrophil subsets play dual roles in tuberculosis by producing inflammasome dependent-IL-1β or suppressing T-cells via PD-L1": sup figures

### Slide 1
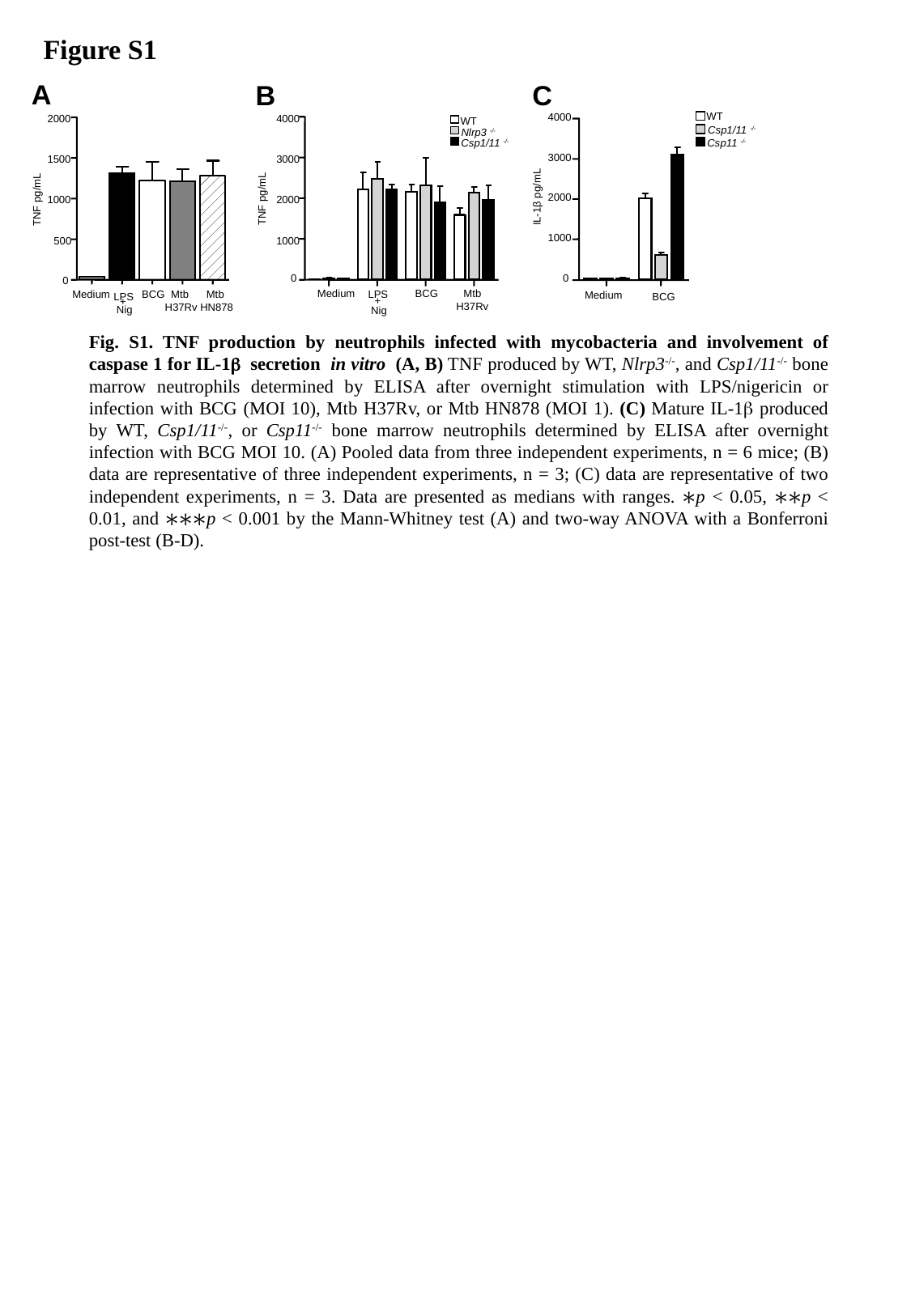

#### Figure S1
A
C
B
WT
Csp1/11 ^-/-^
Csp11 ^-/-^
4000
3000
IL-1β pg/mL
2000
1000
0
Medium
BCG
4000
WT
Nlrp3 ^-/-^
Csp1/11 ^-/-^
3000
2000
1000
0
Medium
BCG
Mtb
H37Rv
LPS
+
Nig
2000
1500
TNF pg/mL
TNF pg/mL
1000
500
0
Medium
BCG
Mtb
 H37Rv
Mtb
HN878
LPS
+
Nig
	Fig. S1. TNF production by neutrophils infected with mycobacteria and involvement of caspase 1 for IL-1β secretion in vitro (A, B) TNF produced by WT, Nlrp3^-/-^, and Csp1/11^-/-^ bone marrow neutrophils determined by ELISA after overnight stimulation with LPS/nigericin or infection with BCG (MOI 10), Mtb H37Rv, or Mtb HN878 (MOI 1). (C) Mature IL-1β produced by WT, Csp1/11^-/-^, or Csp11^-/-^ bone marrow neutrophils determined by ELISA after overnight infection with BCG MOI 10. (A) Pooled data from three independent experiments, n = 6 mice; (B) data are representative of three independent experiments, n = 3; (C) data are representative of two independent experiments, n = 3. Data are presented as medians with ranges. ∗p < 0.05, ∗∗p < 0.01, and ∗∗∗p < 0.001 by the Mann-Whitney test (A) and two-way ANOVA with a Bonferroni post-test (B-D).

### Slide 2
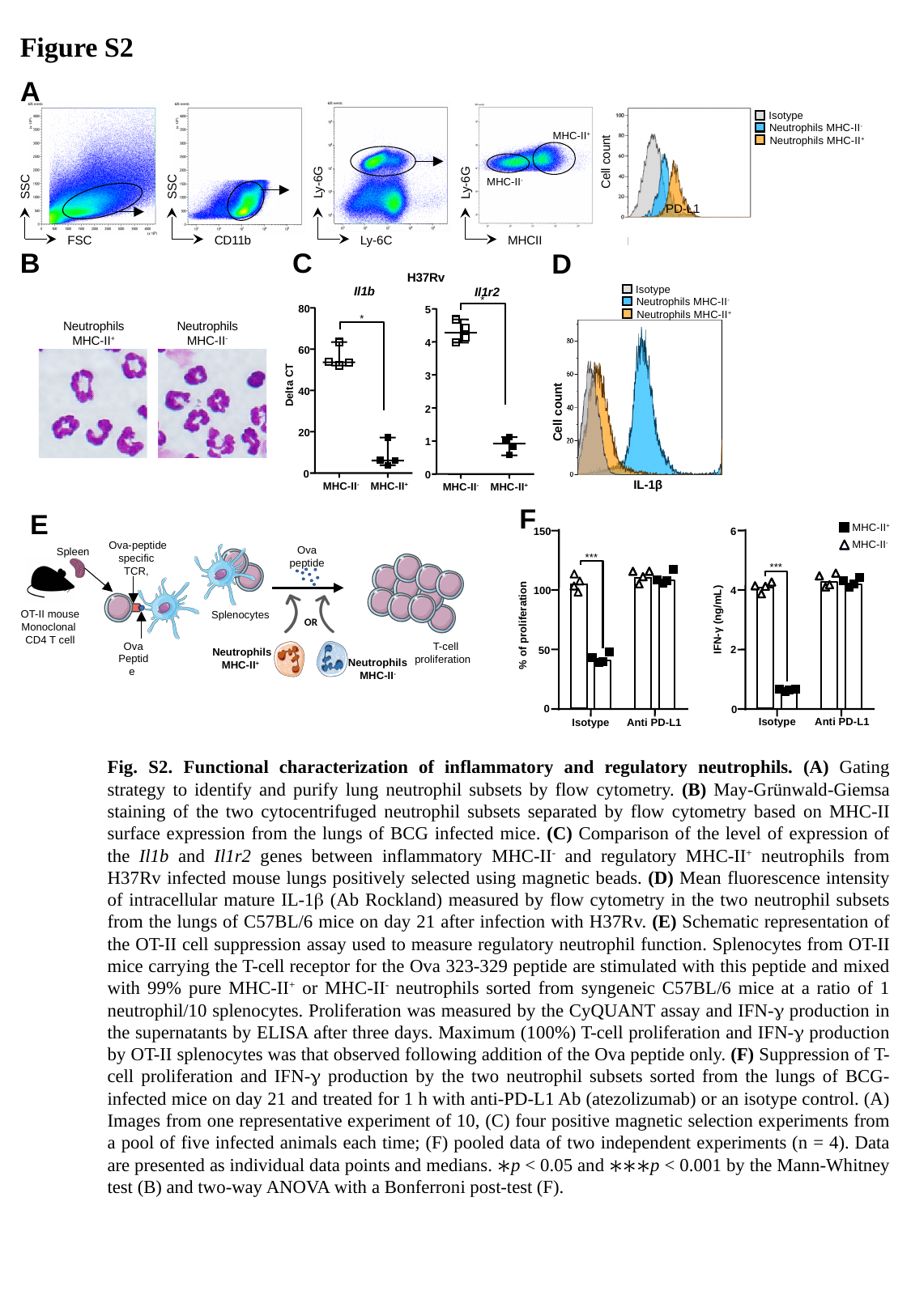

Figure S2
A
Ly-6G
Ly-6C
SSC
FSC
SSC
CD11b
Ly-6G
MHCII
Isotype
Neutrophils MHC-II^-^
Neutrophils MHC-II^+^
 MHC-II^+^
Cell count
MHC-II^-^
PD-L1
B
C
D
H37Rv
Il1b
Il1r2
*
80
5
*
4
60
3
Delta CT
40
2
20
1
0
0
MHC-II^-^
MHC-II^+^
MHC-II^+^
MHC-II^-^
Isotype
Neutrophils MHC-II^-^
Neutrophils MHC-II^+^
 IL-1β
Cell count
Neutrophils MHC-II^+^
Neutrophils MHC-II^-^
F
MHC-II^+^
MHC-II^-^
6
150
***
***
100
4
IFN-γ (ng/mL)
% of proliferation
2
50
0
0
Isotype
Anti PD-L1
Anti PD-L1
Isotype
E
Ova-peptide specific TCR,
OT-II mouse
Monoclonal
CD4 T cell
Ova Peptide
Ova peptide
Spleen
Splenocytes
Neutrophils
MHC-II^+^
 T-cell proliferation
OR
Neutrophils
MHC-II^-^
	Fig. S2. Functional characterization of inflammatory and regulatory neutrophils. (A) Gating strategy to identify and purify lung neutrophil subsets by flow cytometry. (B) May-Grünwald-Giemsa staining of the two cytocentrifuged neutrophil subsets separated by flow cytometry based on MHC-II surface expression from the lungs of BCG infected mice. (C) Comparison of the level of expression of the Il1b and Il1r2 genes between inflammatory MHC-II^-^ and regulatory MHC-II^+^ neutrophils from H37Rv infected mouse lungs positively selected using magnetic beads. (D) Mean fluorescence intensity of intracellular mature IL-1β (Ab Rockland) measured by flow cytometry in the two neutrophil subsets from the lungs of C57BL/6 mice on day 21 after infection with H37Rv. (E) Schematic representation of the OT-II cell suppression assay used to measure regulatory neutrophil function. Splenocytes from OT-II mice carrying the T-cell receptor for the Ova 323-329 peptide are stimulated with this peptide and mixed with 99% pure MHC-II^+^ or MHC-II^-^ neutrophils sorted from syngeneic C57BL/6 mice at a ratio of 1 neutrophil/10 splenocytes. Proliferation was measured by the CyQUANT assay and IFN-γ production in the supernatants by ELISA after three days. Maximum (100%) T-cell proliferation and IFN-γ production by OT-II splenocytes was that observed following addition of the Ova peptide only. (F) Suppression of T-cell proliferation and IFN-γ production by the two neutrophil subsets sorted from the lungs of BCG-infected mice on day 21 and treated for 1 h with anti-PD-L1 Ab (atezolizumab) or an isotype control. (A) Images from one representative experiment of 10, (C) four positive magnetic selection experiments from a pool of five infected animals each time; (F) pooled data of two independent experiments (n = 4). Data are presented as individual data points and medians. ∗p < 0.05 and ∗∗∗p < 0.001 by the Mann-Whitney test (B) and two-way ANOVA with a Bonferroni post-test (F).

### Slide 3
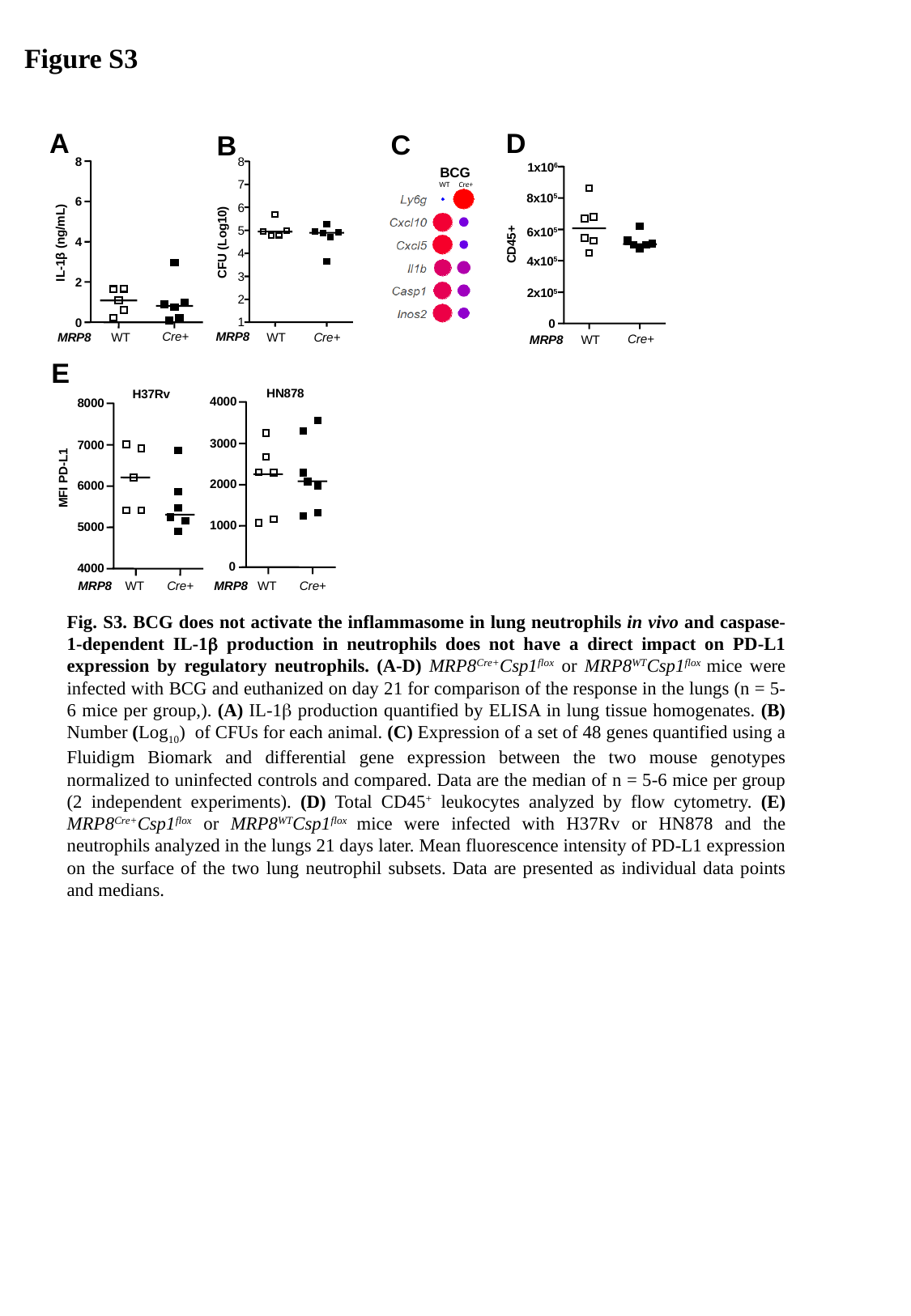

Figure S3
A
D
C
B
8
8
BCG
Cre+
WT
1x10^6^
7
8x10^5^
6
6
5
6x10^5^
4
CFU (Log10)
IL-1β (ng/mL)
CD45+
4
4x10^5^
3
2
2x10^5^
2
1
0
0
Cre+
MRP8
Cre+
MRP8
WT
WT
Cre+
MRP8
WT
E
HN878
H37Rv
4000
8000
3000
7000
MFI PD-L1
2000
6000
1000
5000
0
4000
MRP8
MRP8
WT
Cre+
WT
Cre+
	Fig. S3. BCG does not activate the inflammasome in lung neutrophils in vivo and caspase-1-dependent IL-1β production in neutrophils does not have a direct impact on PD-L1 expression by regulatory neutrophils. (A-D) MRP8^Cre+^Csp1^flox^ or MRP8^WT^Csp1^flox^ mice were infected with BCG and euthanized on day 21 for comparison of the response in the lungs (n = 5-6 mice per group,). (A) IL-1β production quantified by ELISA in lung tissue homogenates. (B) Number (Log_10_) of CFUs for each animal. (C) Expression of a set of 48 genes quantified using a Fluidigm Biomark and differential gene expression between the two mouse genotypes normalized to uninfected controls and compared. Data are the median of n = 5-6 mice per group (2 independent experiments). (D) Total CD45^+^ leukocytes analyzed by flow cytometry. (E) MRP8^Cre+^Csp1^flox^ or MRP8^WT^Csp1^flox^ mice were infected with H37Rv or HN878 and the neutrophils analyzed in the lungs 21 days later. Mean fluorescence intensity of PD-L1 expression on the surface of the two lung neutrophil subsets. Data are presented as individual data points and medians.

### Slide 4
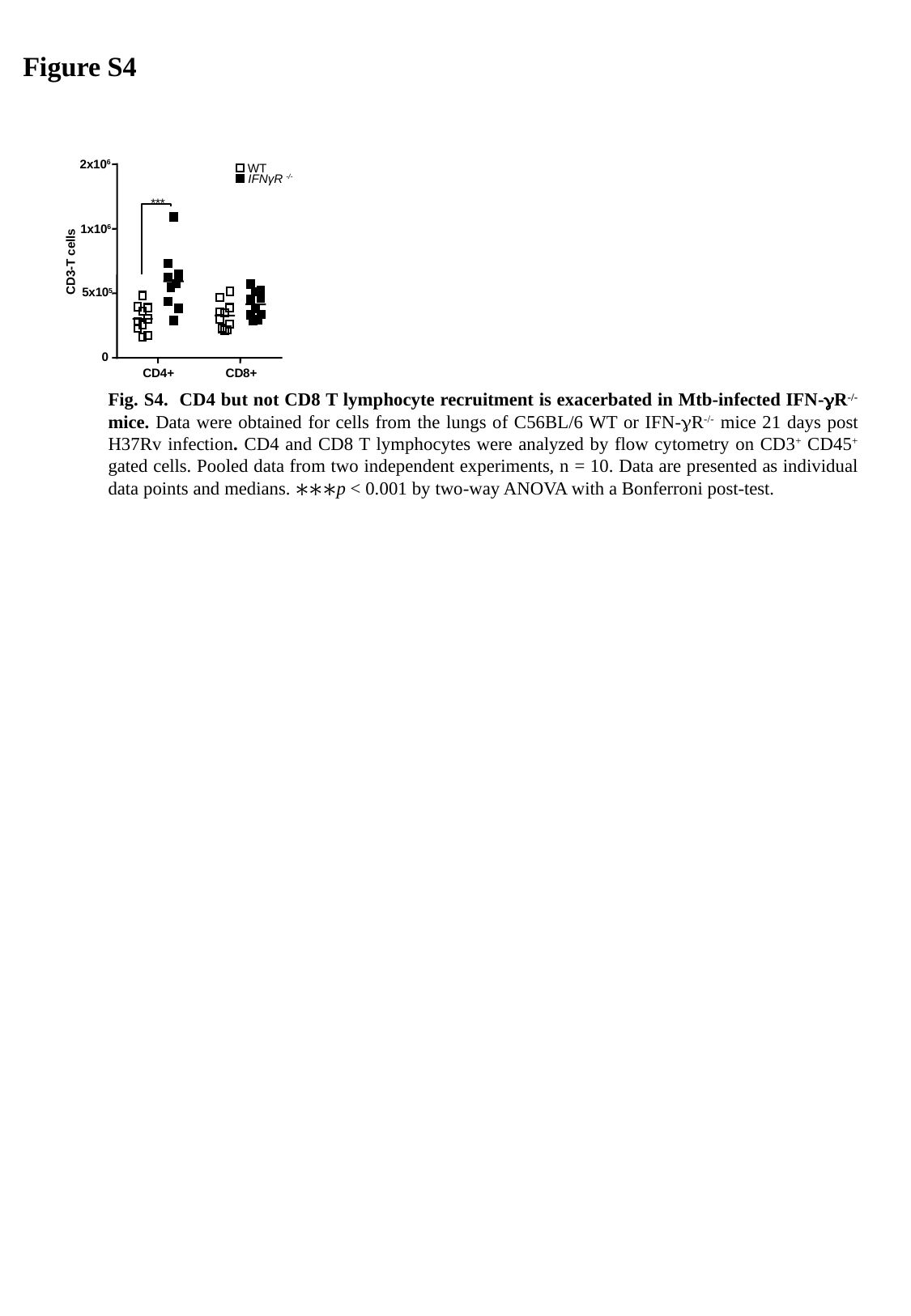

Figure S4
2x10^6^
WT
IFNγR ^-/-^
***
1x10^6^
CD3-T cells
5x10^5^
0
CD4+
CD8+
	Fig. S4. CD4 but not CD8 T lymphocyte recruitment is exacerbated in Mtb-infected IFN-γR^-/-^ mice. Data were obtained for cells from the lungs of C56BL/6 WT or IFN-γR^-/-^ mice 21 days post H37Rv infection. CD4 and CD8 T lymphocytes were analyzed by flow cytometry on CD3^+^ CD45^+^ gated cells. Pooled data from two independent experiments, n = 10. Data are presented as individual data points and medians. ∗∗∗p < 0.001 by two-way ANOVA with a Bonferroni post-test.

### Slide 5
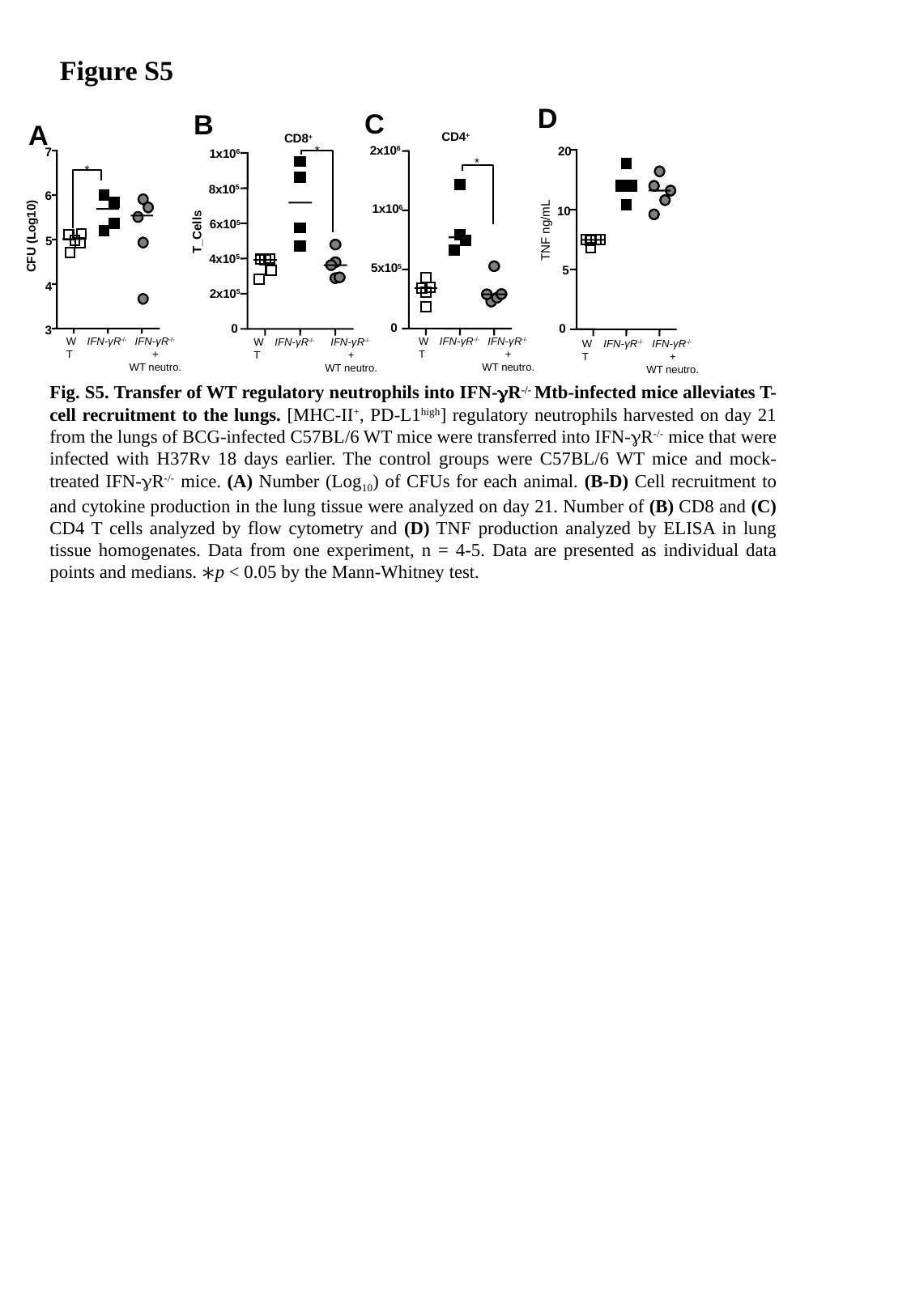

Figure S5
D
20
10
5
0
TNF ng/mL
IFN-γR^-/-^
IFN-γR^-/-^
+
WT neutro.
WT
C
B
A
7
*
6
CFU (Log10)
5
4
3
IFN-γR^-/-^
IFN-γR^-/-^
+
WT neutro.
WT
CD4^+^
CD8^+^
*
1x10^6^
8x10^5^
6x10^5^
4x10^5^
2x10^5^
0
2x10^6^
*
1x10^6^
5x10^5^
0
T_Cells
IFN-γR^-/-^
IFN-γR^-/-^
+
WT neutro.
WT
IFN-γR^-/-^
IFN-γR^-/-^
+
WT neutro.
WT
	Fig. S5. Transfer of WT regulatory neutrophils into IFN-γR^-/-^ Mtb-infected mice alleviates T-cell recruitment to the lungs. [MHC-II^+^, PD-L1^high^] regulatory neutrophils harvested on day 21 from the lungs of BCG-infected C57BL/6 WT mice were transferred into IFN-γR^-/-^ mice that were infected with H37Rv 18 days earlier. The control groups were C57BL/6 WT mice and mock-treated IFN-γR^-/-^ mice. (A) Number (Log_10_) of CFUs for each animal. (B-D) Cell recruitment to and cytokine production in the lung tissue were analyzed on day 21. Number of (B) CD8 and (C) CD4 T cells analyzed by flow cytometry and (D) TNF production analyzed by ELISA in lung tissue homogenates. Data from one experiment, n = 4-5. Data are presented as individual data points and medians. ∗p < 0.05 by the Mann-Whitney test.
