## Supplementary material for "Neutrophil subsets play dual roles in tuberculosis by producing inflammasome dependent-IL-1β or suppressing T-cells via PD-L1": sup material

**Fig. S5.** **Transfer of WT regulatory neutrophils into IFN-γR^-/-^ Mtb-infected mice alleviates T-cell recruitment to the lungs.** [MHC-II^+^, PD-L1^high^] regulatory neutrophils harvested on day 21 from the lungs of BCG-infected C57BL/6 WT mice were transferred into IFN-γR^-/-^ mice that were infected with H37Rv 18 days earlier. The control groups were C57BL/6 WT mice and mock-treated IFN-γR^-/-^ mice. **(A)** Number (Log_10_) of CFUs for each animal. **(B-D)** Cell recruitment to and cytokine production in the lung tissue were analyzed on day 21. Number of **(B)** CD8 and **(C)** CD4 T cells analyzed by flow cytometry and **(D)** TNF production analyzed by ELISA in lung tissue homogenates. Data from one experiment, n = 4-5. Data are presented as individual data points and medians. ∗*p* < 0.05 by the Mann-Whitney test.

**Supplementary Tables**

| **Target** | **Clone** | **Supplier** | **Isotype** | **Dye** |
| --- | --- | --- | --- | --- |
| CD45 | 30-F11 | BioLegend | Rat IgG_2b, κ_ | BV605 |
| CD45 | 30-F11 | BD Biosciences | Rat IgG_2b, κ_ | PerCP-Cy5.5 |
| CD11b | M1/70 | BD Biosciences | Rat IgG_2b, κ_ | V450 |
| CD11c | REA754 | Miltenyi | REAffinity | APC |
| Ly-6G | 1A8 | BD Biosciences | Rat IgG_2a, κ_ | PE |
| Ly-6C | AL-21 | BD Biosciences | Rat IgM | APC |
| MHC-II | 2G9 | BD Biosciences | Rat IgG_2a, κ_ | FITC |
| MHC-II | 2G9 | BD Biosciences | Rat IgG_2a, κ_ | PE |
| CD274(PD-L1) | MIH5 | BD Biosciences | Rat IgG_2a, I_ | BV711 |
| CD3 | 17A2 | BioLegend | Rat IgG_2b, κ_ | PerCP-Cy5.5 |
| CD4 | RM4-4 | BioLegend | Rat IgG_2b, κ_ | FITC |
| CD8 | 53-6.7 | BioLegend | Rat IgG_2a, κ_ | BV510 |
| Mature IL-1β | - | Rockland | Rabbit IgG | Biotin |
| Cleaved IL-1 β | E7V2A | Cell Signaling technology | Rabbit IgG | - |
| Pro- IL-1β | D3H1Z | Cell Signaling technology | Rabbit IgG | - |
| GAPDH | D16H11 | Cell Signaling technology | Rabbit IgG | - |
| Streptavidin | - | BD Biosciences | - | FITC |
| Viability dye | - | eBiosciences | - | eFluor 780 |
| Anti-Rabbit IgG | - | Cell Signaling technology | Goat IgG | HRP |

Table S1. list of antibodies used in this study

| **Gene** | **Forward** | **Reverse** |
| --- | --- | --- |
| *Adgre1* | CAGCCAGAGGGGACAAGATGA | TCCTGGGGCCCCTGTAGAT |
| *Aim2* | TGGGTGGCGTCAGGAAGTTT | GGCCGGTCAACAACAGCATTT |
| *Arg1* | TAGAGAAAGGCCCTGCAGCA | TCAACAAAGGCCAGGTCCCC |
| *Atg7* | CCTGACCTTCGCGGACCTAA | GCCTTCGGCTCGACACAGAT |
| *Casp1* | GGCATGCCGTGGAGAGAAAC | TGGGCCTTCTTAATGCCATCATC |
| *Casp3* | GAGCTTGGAACGGTACGCTAAG | GTCCACTGACTTGCTCCCATGTA |
| *Casp4* | TCCCAGATGCCCACCATTGA | TCAGTTGCTTGTTGCTTTGTTCTC |
| *Casp8* | CCACAGACCCCAGACAGAGAA | AGAGGTAGAAGAGCTGTAACCTTAT |
| *Cat* | AGATTGCCTTCTCCGGGTGG | CCCGACTGTCCGACATGGTG |
| *Ccl20* | CTTCCTTCCAGAGCTATTGTGG | TCATCCATTGGACAAGTCCACTG |
| *Ccl5* | TCTCTGCAGCTGCCCTCACC | TCTTGAACCCACTTCTTCTC |
| *Ccl7* | CCTGGGAAGCTGTTATCTTCAA | CCTCCTCGACCCACTTCTGAT |
| *Ccl8* | ACGCTAGCCTTCACTCCAAAATC | GGCTGACAGGGACAGCTATGA |
| *Ccr1* | CTCTGGAAACACAGACTCACTGTC | TTGGCATGGAGTGGAGTCCC |
| *Cd14* | AAGCCCGTGGAACCTGGAAG | CACGCTCCATGGTCGGTAGA |
| *Cd209a* | ACAGTCAAGTCCCCTTGGCA | AGTCGATCTACGCCAGCCTTC |
| *Cd28* | TGGCTCTTTGTGTTATCTGGACAAA | GGGCGTAGGGCTGGTAAGG |
| *Cd40* | TCACCATTTTCGGGGTGTTTCT | CCGCAGGGGGTAAGATCTCATT |
| *Cd74* | TGGATGGCGTGAACTGGAAGA | TCCTGGCACTTGGTCAGTACTTT |
| *Cd80* | CGACTCGCAACCACACCATTA | GGAGGGTCTTCTGGGGGTTTT |
| *Cd86* | CGCGTAAGAGTGGCTCCTGTA | CCAAGCCCATGGTGCATCT |
| *Mb21d1 (Cgas)* | AAGGCTGGCTGGGCACAAA | CGCCAGGTCTCTCCTTGAAAA |
| *Clec7a* | GGGGATCAGAGAAAGGAAGCCA | CAGCACTGCAGCAACCACTAC |
| *Col1a1* | GCTACTACCGGGCCGATGAT | CGATCCAGTACTCTCCGCTCTT |
| *Col3a1* | ACCCAGAGGAGTAGCTGGAGAA | TCCGGGCATACCCCGTATC |
| *Ctla4* | GTCTGTGTGGGTTCAAACACATCT | AGGTCCTCAGGGAGCAGAGTAA |
| *Ctsg* | AATGTGCGCCAATCGCTTCC | GAATCACCCCTGAAGGCAGAC |
| *Ctsk* | CCCATATGTGGGCCAGGATGAAA | TTCTCGTTCCCCACAGGAATCT |
| *Cxcl1* | CGCTCGCTTCTCTGTGCAGC | GTGGCTATGACTTCGGTTTGG |
| *Cxcl10* | CACGTGTTGAGATCATTGCCA | GCGTGGCTTCACTCCAGTTA |
| *Cxcl2* | GCTGCTGGCCACCAACCACC | TGAGAGTGGCTATGACTTCTG |
| *Cxcl5* | CCCTACGGTGGAAGTCATAGCTAAA | GCCGTTCTTTCCACTGCGAG |
| *Cxcr1* | CCAGCTGGTGCCTCAGATCAAA | TGGGCAGCATTCCCGTGATA |
| *Cxcr2* | TCAACCAGCCCTGACAGCTC | ACTTAATCCTGCAGTAGTTCTACGA |
| *Cxcr3* | CCAAGCCATGTACCTTGAGGTTAGT | AGTCGCTCTCGTTTTCCCCATA |
| *Cybb* | TGGGATGAATCTCAGGCCAATCA | CCAGTTGGGCCGTCCATACA |
| *Ela2* | TCAGCAGCCCACTGTGTGAA | AGAAGGTCTGTCGAGTGCGC |
| *Fcgr1* | TGATTCTTACCAGCTTTGGAGATGA | CCACCGACTGGAACCCAAAG |
| *Gmcsf* | TGCAGACCCGCCTGAAGATA | GGCCTGGGCTTCCTCATTTT |
| *Gsdmd* | TCCCGGGTTGAGCAGACAAT | CGATGGCATGGTCCTCGATTT |
| *Hprt1* | CAGTCCCAGCGTCGTGATTA | TGGCCTCCCATCTCCTTCAT |
| *Ifi204* | GCTGATTCTGGATTGGGCAAACT | CAGTGATGTTTCTCCTGTTACTTCT |
| *Ifna* | CTCCTAGACTCATTCTGCAATGA | GGGCTCTCCAGACTTCTGCTCTG |
| *Ifnar1* | CTCCCCGCAGTATTGATGAGTTTT | CTCAGGCGCGTGCTTTACTT |
| *Ifnar2* | CACCGTCTGCTTTTGATGGGTAT | GGTGGGCCAGACTTGTTCTC |
| *Ifnb1* | AAGCAGCTCCAGCTCCAAGAA | TGGATGGCAAAGGCAGTGTAAC |
| *Ifng* | TCTTCTTGGATATCTGGAGGAA | AGCTCATTGAATGCTTGGCGCTG |
| *Ifngr1* | GGTGCCTGTACCGACGAATG | GGTGCCTGTACCGACGAATG |
| *Ifnlr1* | GGTGCCTGTACCGACGAATG | CAGTCCAGGAACCCGAATACAC |
| *Il10* | ATGCTGCCTGCTCTTACTGAC | CTGGGGCATCACTTCTACCAG |
| *Il10rb* | CGAGCCCGCAGCTGTTT | GATCTTGGAAAGACCTGTAACTTTC |
| *IL12p40* | CTCACATCTGCTGCTCCACAA | GACGCCATTCCACATGTCACT |
| *Il17* | TCCAGAAGGCCCTCAGACTA | AGCATCTTCTCGACCCTGAA |
| *Il18* | CCTCTTGGCCCAGGAACAATG | ACAGTGAAGTCGGCCAAAGTT |
| *Il18r1* | CACAACGATCCTGAAAACAAGAGAT | AAGGTTCTCCCTCTACCACATGAA |
| *Il18rap* | CCAGTCTCAGCTGCCAAAGT | AGGAAGTAGTCTTCCATCCTTGTA |
| *Il1a* | CGCTTGAGTCGGCAAAGAAATCA | TGCAAGTCTCATGAAGTGAGCCA |
| *Il1b* | TCTAATGCCTTCCCCAGGGC | GACCTGTCTTGGCCGAGGAC |
| *Il1r1* | ACCGTGAACAACACAAATGGAGAA | ATGGTGTCGCCGTGCATTTT |
| *Il1r2* | GGAGACCCCACACGCCTATT | GGGTTCCGTGGTTGTTCCTTTG |
| *Il1rn* | GGAAGACCTTGTGTCCTGTTTAG | GGCACCATGTCTATCTTTTCTTC |
| *Il33* | GCTGCGTCTGTTGACACATT | GACTTGCAGGACAGGGAGAC |
| *Il4* | ACGGAGATGGATGTGCCAAACGTC | AACTTTCCAGGAAGTCTTTCAG |
| *Il6* | GAGGATACCACTCCCAACAGACC | AAGTGCATCATCGTTGTTCATACA |
| *Il6ra* | CAACACCACCAACGGGAAGA | GGATCCGGCTGCACCATTTTT |
| *Inos2* | GCCACCTTGGTGAAGGGACT | ACGTTCTCCGTTCTCTTGCAGT |
| *Irf1* | CCAGCATCTCGGGCATCTTT | GAGTGATTGGCATGGTGGCTTTG |
| *Irf3* | CAATTCCTCCCCTGGCTAGA | AGATGCCAAAGTCAGCCATCTG |
| *Irf7* | GGTCCAGCGAGTGCTGTTTG | CACAGCCCAGGCCTTGAAGA |
| *Irf9* | TCAGGCCCTGCCCATTTCTTC | TTTGCCTGAGGCCATCCTTCT |
| *Itgam* | GCTCTCATCACTGCTGGCCT | GTTACTGAGGTGGGGCGTCT |
| *Ly6g* | CCCTGCTGTATAGGCACCCC | ATGCCTCCAGGGTCAAGAGC |
| *MhcII* | GGAGCAAGATGTTGAGCGGC | GCCTCGAGGTCCTTTCTGAC |
| *Mki67* | CATCATTGACCGCTCCTTTAGGTAT | GGTATCTTGACCTTCCCCATCAG |
| *Mlkl* | GTCTTTCTGGCAGAGAACGAATCT | TACACCTTCTTGTCCGTGGATTCT |
| *Mmp7* | TTTGATGGGCCAGGGAACACTCTA | ATGGGTGGCAGCAAACAGGAAGT |
| *Mmp8* | ACAGGGAACCCAGCACCTATT | TGGGGTTGTCTGAAGGTCCATAG |
| *MMP9* | ACCACCACCACCACACACAA | CTGCCTCCACTCCTTCCCAG |
| *Mpo* | TGTCCGTGTCAAGTGGCTGT | GGGGCTTCGTCTGTTGTTGC |
| *Myd88* | CACTCGCAGTTTGTTGGATG | CGCAGGATACTGGGAAAGTC |
| *Nfkb* | CCACGAGGCAGCACATAGATGA | GCAGTGGGCTGTCTCCAGTA |
| *Nlrc4* | GACGCTTTGACTCACCACAATGAA | CTGCTCCAAGGGCTCACAGTA |
| *Nlrp3* | TGCGTGTTCTCTGTATACCACATCT | GGGCTTAGGTCCACACAGAAAGT |
| *Oas1* | GGTCAAGGGCAAAGGCACCA | TCTCATGCTGAACCTCGCACA |
| *Oasl2* | CCCACCAACAACCTGGGAAGA | ACATCCCTCGCTCGCTGTA |
| *Orl1* | GGCTGAGGTCCTCGACTGTTTC | GGAAGAAAGCAAATGCAGACCTTTA |
| *P2rx7* | CGGATCCAGAGCACGAATTATGG | CGCTCACCAAAGCAAAGCTAATGT |
| *Pdcd1* | CTGGAGCAGAGCTCGTGGTAA | AGCTCCTCTGGCCTCTGACATA |
| *Pdl1* | GCAGGCGTTTACTGCTGCAT | TGCGGTATGGGGCATTGACT |
| *Ppia* | GCTGGACCAAACACAAACGG | CCAAAGACCACATGCTTGCC |
| *Prtn3* | AGCAGGCATATGCTTCGGAGA | CCCCGCAGCACGTTTTGAAT |
| *Ptgs2* | AGCCAGGCAGCAAATCCTTG | ACTGTGTTTGGGGTGGGCTT |
| *Rpl4* | GACCAGTGCTGAGTCTTGGG | GTATTCACTCTGCGGTGCCA |
| *S100a8* | TCCTTTGTAAGCTCCGTCTTC | CTTCTCCAGTTGAGACGGCA |
| *S100a9* | GTGGAAGCACAGTTGGCAAC | TGGGTTGTTCTCATGCAGCT |
| *Slc11a1* | TGGCCATTGGGGCTCAGAT | TCCAGCTTGCGCAAACCATAG |
| *Socs1* | GACGCCTGCGGCTTCTATTG | GACTGTCGCGCACCAAGAA |
| *Socs2* | CTGGAGCCTCCGGGAATG | TCCCCAGTACCATCCTGTTTGA |
| *Socs3* | CGCGGGCACCTTTCTTATCC | GGGTCACTCTGCAGCGAAAA |
| *Sod2* | TGAGCCCTAAGGGTGGTGGA | ACGGCTGTCAGCTTCTCCTT |
| *Stat1* | TCAAGCTGAGACTGTTGGTGA | TGTGTGCGTACCCAAGATGT |
| *Stat2* | AGCATTTGGCTACCTGGATTGA | GCCATTGGGAAAGGTCTGAAT |
| *Stat3* | GCTGCCCCGTACCTGAAGA | TGTCAAACGTGAGCGACTCAAA |
| *Tbk1* | TCAGGCTGGCCACCAGAAA | TCTCTTGGATGCGTGCCTTCT |
| *Timp1* | TCCTAGAGACACACCAGAGCAGAT | GGGAACCCATGAATTTAGCCCTTAT |
| *Tlr2* | GCATCCGAATTGCATCACCG | CATCACACACCCCAGAAGCA |
| *Tlr4* | TTATCCAGGTGTGAAATTGAAAC | GCCACATTGAGTTTCTTTAAGG |
| *Tmem173 (sting)* | GGTCTAGGAAGCAGAAGATGCCATA | TCAGGCTGGCCACCAGAAA |
| *Tnfa* | ATGAGCACAGAAAGCATGATC | TACAGCCTTGTCACTCGAATT |
| *Zpb1* | CGCCAAGGCTCTGGGAATGA | TGTGTGACTCCAGAATGAGCTATGT |

Table S2: list of primers used for transcriptomic analysis (qRT-PCR and medium throughput)
